# Single cell spatial transcriptomic profiling of childhood-onset lupus nephritis reveals complex interactions between kidney stroma and infiltrating immune cells

**DOI:** 10.1101/2023.11.09.566503

**Authors:** Patrick Danaher, Nicholas Hasle, Elizabeth D. Nguyen, Kristen Hayward, Natalie Rosenwasser, Charles E. Alpers, Robyn C. Reed, Daryl M. Okamura, Sarah K. Baxter, Shaun W. Jackson

## Abstract

Children with systemic lupus erythematosus (SLE) are at increased risk of developing kidney disease, termed childhood-onset lupus nephritis (cLN). Single cell transcriptomics of dissociated kidney tissue has advanced our understanding of LN pathogenesis, but loss of spatial resolution prevents interrogation of in situ cellular interactions. Using a technical advance in spatial transcriptomics, we generated a spatially resolved, single cell resolution atlas of kidney tissue (>400,000 cells) from eight cLN patients and two controls. Annotated cells were assigned to 35 reference cell types, including major kidney subsets and infiltrating immune cells. Analysis of spatial distribution demonstrated that individual immune lineages localize to specific regions in cLN kidneys, including myeloid cells trafficking to inflamed glomeruli and B cells clustering within tubulointerstitial immune hotspots. Notably, gene expression varied as a function of tissue location, demonstrating how incorporation of spatial data can provide new insights into the immunopathogenesis of SLE. Alterations in immune phenotypes were accompanied by parallel changes in gene expression by resident kidney stromal cells. However, there was little correlation between histologic scoring of cLN disease activity and glomerular cell transcriptional signatures at the level of individual glomeruli. Finally, we identified modules of spatially-correlated gene expression with predicted roles in induction of inflammation and the development of tubulointerstitial fibrosis. In summary, single cell spatial transcriptomics allows unprecedented insights into the molecular heterogeneity of cLN, paving the way towards more targeted and personalized treatment approaches.

## Introduction

Childhood-onset systemic lupus erythematosus (cSLE) is characterized by increased incidence of kidney involvement, termed lupus nephritis (LN). As with adult SLE, the failure to achieve clinical remission in lupus nephritis is associated with long-term kidney damage. This is particularly concerning in children who have the potential to accrue kidney damage over decades (*1*). Unfortunately, despite recent treatment advances, less than half of all patients in large clinical trials achieve complete renal response (*2–4*), underscoring the need to both develop additional therapies targeted against the underlying mechanisms and to resolve the heterogeneity of disease in a manner that allows precision management.

One strategy to interrogate the heterogeneity of human lupus nephritis is the use of single-cell transcriptomics (scRNA-Seq), a powerful tool allowing a complete catalogue of cell types and cell states in normal and diseased tissues. In 2019, the Accelerating Medicines Partnership (AMP) in SLE network published two studies describing the scRNA-Seq landscape of infiltrating immune cells (*5*) and renal tubular stomal cells (*6*) in human lupus nephritis. Despite this major advance in uncovering the molecular heterogeneity of lupus nephritis, there were several limitations to these data. First, tissue dissociation required for scRNA-seq analysis eliminates all spatial information; a limitation exacerbated by the separate analysis of immune and stromal compartments in the AMP studies. Second, several rare but important cell types (such as podocytes and mesangial cells) were not captured in these earlier studies. Additionally, kidney tissue dissociation can induce an artifactual stress response (*7*) and the need for central processing of fresh kidney biopsy tissue limits sample availability (*8*).

In this study, we leverage a technical advance in spatial transcriptomics to study childhood-onset lupus nephritis (cLN) at single cell resolution. The CosMx Spatial Molecular Imager (SMI) is a new platform allowing the detection of up to 1000 genes at single (sub)-cellular resolution on archived clinical samples stored as formalin-fixed, paraffin-embedded (FFPE) tissue (*9*). Here, we compare the transcriptional signatures of both infiltrating immune cells and resident kidney stromal cells in cLN. We focused on pediatric subjects since childhood-onset SLE is relatively understudied and pediatric patients are frequently excluded from LN clinical trials. In addition, children typically lack concomitant chronic diseases such as diabetes or long-standing hypertension, allowing a more focused assessment of LN-specific changes.

Using this spatially-resolved atlas of cLN, we demonstrate that resident glomerular cells and infiltrating immune cells in cLN exhibit broad changes in gene expression reflecting both injury responses and initiation of tissue repair mechanisms. In parallel, we demonstrate that individual immune lineages localize to specific regions in cLN kidneys and that their transcriptional signatures vary as a function of tissue location. These data highlight the heterogeneity of human lupus nephritis, both between affected subjects and across individual tissue sections, and emphasize the importance of defining disease-associated transcriptional changes as a function of spatial neighborhoods. Together, our findings provide new insights into the pathogenesis of lupus nephritis, which may inform the development of new therapies targeted at disease mechanisms.

## Results

### Spatially-resolved single cell resolution spatial transcriptomics of childhood-onset lupus nephritis

To discern the spatial heterogeneity of kidney inflammation in cLN, we performed CosMx spatial transcriptomics on diagnostic kidney biopsy tissue from 7 pediatric patients with lupus nephritis (denoted SLE1 to SLE7). FFPE kidney biopsies were collected as part of routine clinical practice and stored prior to CosMx analysis. Except for hydroxychloroquine and corticosteroids, diagnostic kidney biopsies were obtained prior to initiation of immunosuppression, except for subject SLE5 who received a single dose of intravenous cyclophosphamide 10 days prior to biopsy. The cLN cohort included subjects with proliferative (Class III and Class IV) lupus nephritis (*10*). We deliberately selected patients with variable baseline proteinuria (ranging from mild to nephrotic range). We also analyzed 3 serial kidney biopsies from a patient with treatment-resistant Class IV lupus nephritis, obtained over a 16-month period (SLE8). As controls, we included histologically normal kidney tissue from 2 pediatric subjects who underwent nephrectomy for non-inflammatory indications (**Fig. 1A**). The clinical characteristics of the control and cLN cohort subjects are summarized in **Supplemental Tables S1 – S4**.

**Figure 1:**
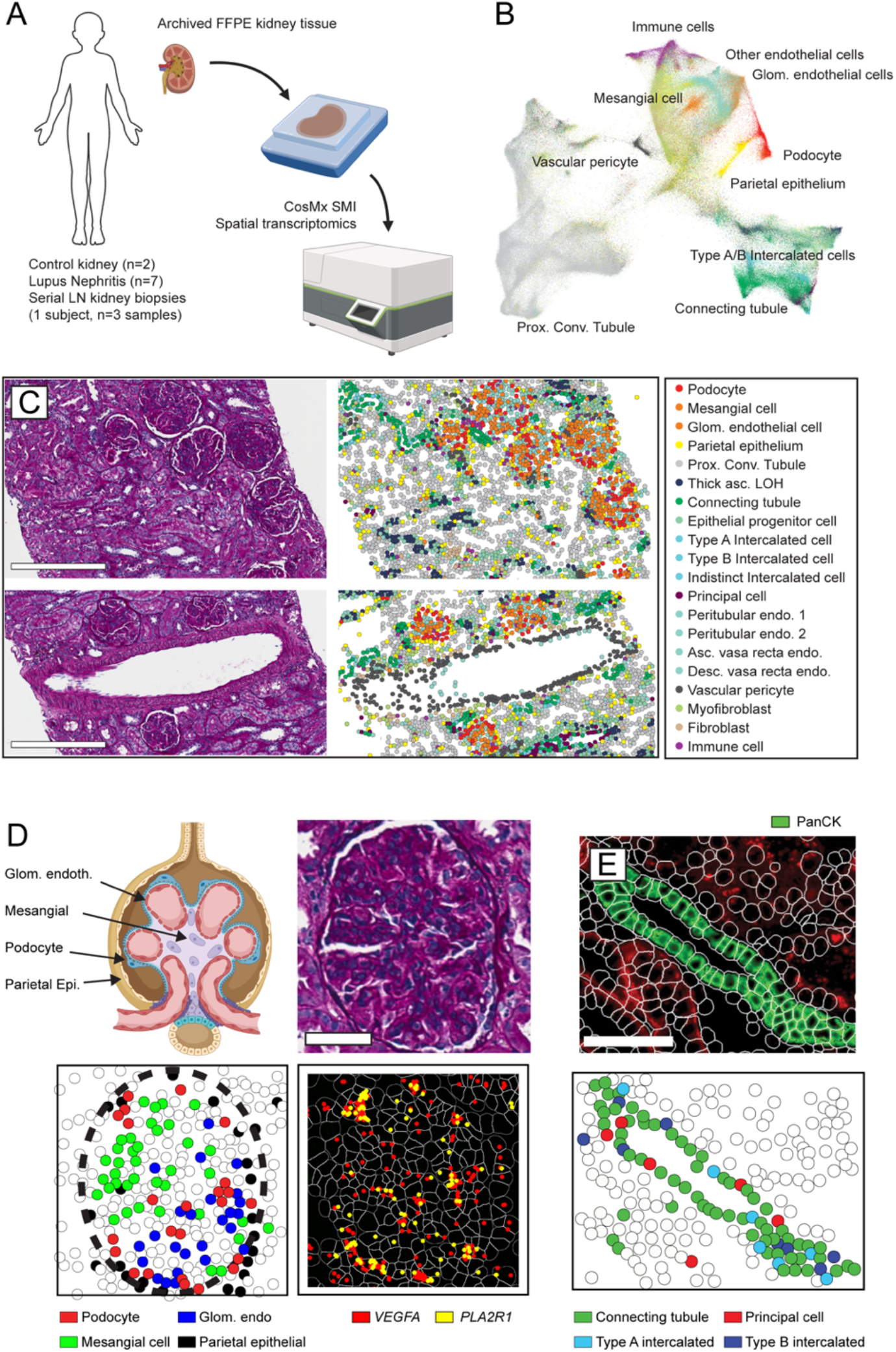
Single cell-resolution spatial transcriptomics in childhood-onset lupus nephritis (cLN). **(A)** Archived kidney biopsy tissue stored in FFPE was profiled using the CosMX Spatial Molecular Imager to obtain single cell-resolution spatial transcriptomic data from 2 healthy controls, 7 subjects with cLN, and 1 subject with treatment-resistant cLN across 3 serial biopsies. **(B)** UMAP projection of kidney and immune cell subsets in combined CosMX dataset. **(C)** Tissue sections from representative cLN subject (SLE6) showing Periodic acid– Schiff (*PAS*)-stained tissue sections (left) compared to equivalent cell types annotated using spatial transcriptomics. Scale bars, 300μm. **(D)** Upper left: schematic of the human glomerulus showing distribution of relevant cell types. PAS-stained histology (upper right) vs. transcriptionally-defined glomerular cell types (lower left) in single glomerulus from representative cLN patient. Lower right: Expression of podocyte-specific genes (*VEGFA*, red; *PLA2R1*, yellow) overlaps with spatial location of podocytes. Scale bars, 50μm. **(E)** Corresponding images showing panCK^+^ regions (upper panel, green) overlapping with transcriptionally-defined Connecting Tubule cells (green), Type A Intercalated cells (light blue), Type B Intercalated cells (dark blue), and Principal cells (red). Scale bars, 100μm.

After data QC and cell segmentation, we identified a total of 447,892 spatially-resolved cells (mean 37,324 cells/biopsy). Since the CosMx platform does not require tissue dissociation, cell dropout was minimal, resulting in a marked increase in annotated cells compared with prior LN scRNA-Seq studies, which annotated an average of ∼100-150 cells per tissue (*5, 6*). The cornerstone of any single cell transcriptomic analysis is cell typing based on mRNA expression. Since CosMx measures a fraction of the total transcriptome (960 genes in this panel) falling within a 4μm tissue section, the resulting data are more sparse than standard scRNA-seq data, rendering established scRNA-seq data analysis tools suboptimal for these data. However, spatial transcriptomics incorporates additional data such as cell imaging, spatial location, and the cell’s “neighborhood”. To incorporate these additional data, we used InSituType, a new tool specifically designed for preprocessing and cell typing with CosMx data, which outperforms established scRNA-seq tools, including Seurat (*11*) and SingleR (*12*). InSituType uses a negative-binomial likelihood model in a naïve Bayes classification scheme to assign individual cells to previously annotated cell types and incorporates the spatial information that is inherent to CosMx data into the posterior using context-specific prior distributions (*13*). We applied this framework for supervised cell typing using the Human Cell Atlas reference dataset (*14*) and a broader set of immune cell profiles (*15*), resulting in individual cells being assigned to 35 reference cell types (**Fig. 1B; Supplemental Figure 1**).

Visualizing spatial relationships together with algorithm-defined cell typing provided robust evidence of the accuracy of cell annotation. By comparing annotated cells with the corresponding Periodic acid–Schiff (PAS)-stained tissue sections, we confirmed that morphologically-defined glomerular, tubulointerstitial, and vascular regions overlapped with anticipated kidney stromal cell populations (**Fig. 1C**). For example, the filtration unit of the kidney, the glomerulus, comprises three major cell types: fenestrated glomerular endothelial cells; mesangial cells; and podocytes. These cell types co-localized into distinct glomerular structures separate from the tubulointerstitium (**Fig. 1D**; **Supplemental Figure 1)**. Notably, fenestrated glomerular endothelial cells could be transcriptionally distinguished from endothelial cells lining the extra-glomerular renal vessels, demonstrating that InSituType is able to discern even closely-related cell subpopulations. Using the BioTuring Lens spatial analysis platform (*16*), we confirmed that *VEGFA* and *PLA2R1* transcript expression mapped to podocytes, in keeping with known roles for these genes in podocyte biology (*17, 18*) (**Fig. 1D**). Finally, as part of the CosMx workflow, tissues are stained with morphology markers, including pan-cytokeratin (panCK), which is known to stain distal tubules of the human kidney. We leveraged this staining pattern to further validate kidney cell typing. Overlaying panCK^+^ immunofluorescence and CosMx cell annotations confirmed localization of connecting tubule cells, type A and B intercalated cells, and principal cells to panCK^+^ distal tubule segments (**Fig. 1E**). In summary, we used CosMx on archived clinical kidney biopsy tissue to generate a spatial transcriptomic atlas that is comprehensive of known kidney cell types.

### Immune landscape of childhood-onset lupus nephritis

Spatial transcriptomics identified 11 immune cell types, producing a granular map of the cLN immune infiltrate. The proportion of immune cells was modestly increased in cLN, albeit with marked variability between cLN subjects (**Fig. 2A**). Macrophages were the most abundant immune lineage representing 73% and 64% of all immune cells in control and cLN biopsies, respectively. However, whereas intrarenal macrophage numbers were similar in cases and controls, the development of cLN was characterized by a relative expansion of specific immune lineages. These included members of the B cell lineage (B cells and plasma cells) and plasmacytoid dendritic cells (pDCs) (**Fig. 2B**), in keeping with known roles for these immune lineages in the pathogenesis of LN (*19–21*).

**Figure 2:**
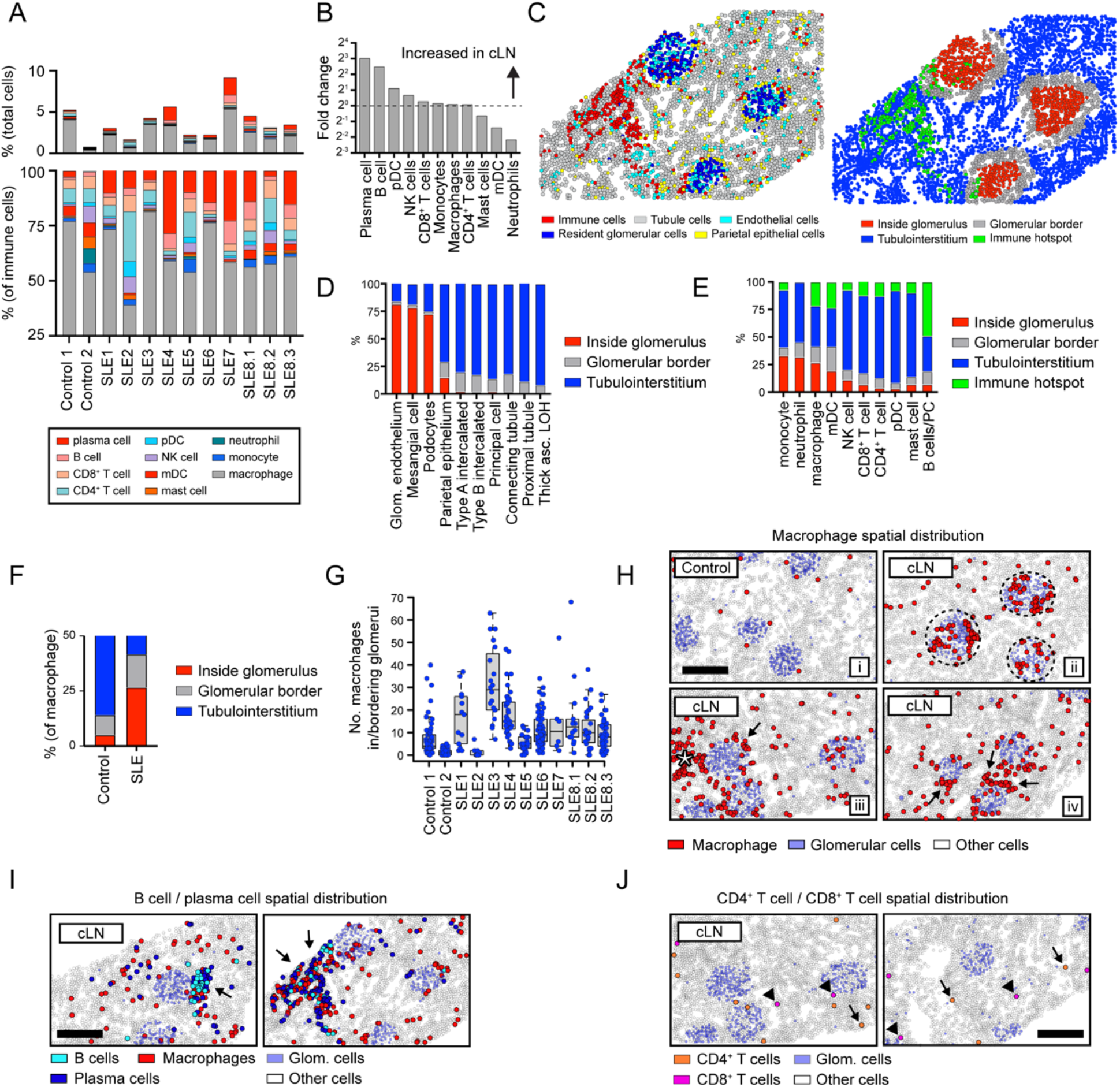
Immune landscape of cLN. **(A)** Immune cell landscape in controls vs. cLN. Upper panel: immune cell infiltrates as % of total cells. Lower panel: relative proportions of immune cell subsets. **(B)** Ratio of immune cell subset percentages in cLN vs controls. **(C)** Corresponding images showing major kidney cell types (left) and computationally-defined anatomical regions (right). Each dot indicates an individual cell colored by cell type (left) or spatial location (right). **(D, E)** Proportion of kidney stromal cells (D) and infiltrating immune cells (E) located within the 4 spatial kidney regions. **(F)** Proportion of macrophages located inside (red) or bordering (grey) glomeruli in controls vs. cLN. **(G)** Total number of macrophages in/bordering glomeruli in individual control and cLN subjects. Each dot equals macrophage number per glomerulus. Boxes show interquartile ranges over each tissue’s glomeruli. **(H)** Representative patterns of macrophage spatial distributions: i) macrophages (red) diffusely distributed in tubulointerstitium in control kidney; ii) cLN macrophages trafficking to glomeruli (dashed circle) in cLN; iii/iv) macrophages surrounding glomeruli (arrows) or clustered within tubulointerstitial immune hotspot (star) in cLN. **(I)** B cells (light blue) and plasma cells (dark blue) colocalize within tubulointerstitial immune hotspots (arrows) in cLN. Macrophages (red) are also enriched within B cell/plasma cell foci. **(J)** Spatial distribution of CD4^+^ T cells (orange; arrows) and CD8^+^ T cells (pink; arrowheads) in cLN. (H-J) Scale bars, 200μm.

Leveraging the spatial information inherent to CosMx data, we examined whether infiltrating immune cells trafficked to distinct anatomical kidney regions in cLN. To do this efficiently, we designed a computational strategy in which tissue sections were partitioned into 4 spatial regions defined as: i) inside glomeruli; ii) bordering glomeruli; iii) within the kidney tubulointerstitium; and i) within “immune hotspots” (regions comprising ≥50 immune cells within 0.5mm of nearest immune neighbor) (**Fig. 2C**). To validate the accuracy of these computationally-defined tissue regions, we quantified the spatial distribution of kidney stromal cells. As predicted, tubular cell subsets uniformly localized to the tubulointerstitium or abutted glomeruli, while ∼70-80% of resident glomerular cells (glomerular endothelial cells, mesangial cells, and podocytes) resided inside glomeruli (**Fig. 2D**).

Incorporating this spatial information into the analysis of cLN immune infiltrates highlighted new features of lupus biology. For example, an increased proportion of myeloid lineages (monocytes, macrophages, neutrophils) trafficked into glomeruli in cLN, while other macrophages surrounded inflamed glomeruli (**Fig. 2E**). Consistent with intra-glomerular myeloid cell infiltration being a characteristic feature of glomerulonephritis, 42% macrophages localized within or bordering glomeruli in cLN compared with 14% in controls (**Fig. 2F**). Although glomerular localization was characteristic of cLN, intra-glomerular macrophage numbers varied markedly between subjects, highlighting the heterogeneity of human SLE (**Fig. 2G**). Of the remaining cLN-associated macrophages residing away from glomeruli, 63% were diffusely distributed in tubulointerstitial regions, while 37% were found in foci of densely clustered immune cells (**Fig. 2H**; star denotes macrophages in tubulointerstitial immune hotspot). Monocytes and monocytic dendritic cells (mDCs) had similar spatial distributions as macrophages, but at much lower frequencies (<2% of SLE sample immune cells; data not shown).

In contrast to macrophages, lymphocytes were infrequent within glomeruli, and instead localized to tubulointerstitial regions. Expanded populations of B cells and plasma cells comprised 21% of all immune cells in cLN. These cells seldom appeared in isolation, with 96% B cells and 94% plasma cells neighboring other immune cells (>1 immune cell within 50 closest neighbors), and ∼50% of B cells/plasma cells residing in tubulointerstitial immune hotspots (**Fig. 2I**). CD4^+^ and CD8^+^ T cells were significantly less abundant than B cells, with only ∼12% localized to immune hotspots and the remainder more diffusely distributed throughout the tubulointerstitium (**Fig. 2J**).

### Macrophage gene expression in cLN varies based on spatial distribution

To further define how spatial context interacts with function, we analyzed immune cell gene expression as a function of tissue distribution. During lupus nephritis pathogenesis, immune-complex deposition promotes recruitment of circulating monocytes to inflamed glomeruli in response to chemotactic signals derived from resident glomerular cells (*22*). Subsequently, these infiltrating monocytes undergo a phenotypic transition, differentiating into activated macrophages with diverse pro-inflammatory and reparative potentials. Based on this model, we quantified differentially expressed genes (DEG) in intra-vs. extra-glomerular macrophages in cLN kidneys.

Notably, intra-glomerular macrophages upregulated genes linked to function, including antigen uptake and lipid metabolism (*FABP4*, *FABP5*, *CD36*), fibroinflammatory pathways (*S100A4* (*23, 24*), *S100A9* (*25*)), cell surface adhesion and migration (*COTL1*, *EMP3*), chemokine production (*CCL3*, *CCL4*), and tissue remodeling (*MMP19*) (**Fig. 3A**). High-magnification images confirmed that macrophages located adjacent to mesangial cells in cLN expressed *FABP4*, *FABP5*, *ITGAX*, *S100A4*, *CD36*, *CD52*, and *COTL1* gene transcripts (**Fig. 3B**). In contrast, intra-glomerular macrophages lacked expression of genes commonly expressed in tissue-resident macrophages (TRM), including *SELENOP*, *MS4A4A*, *MRC1* (encoding CD206), and C1q components (*C1QA*, *C1QB*, *C1QC*) (*26*) (**Fig. 3A**).

**Figure 3:**
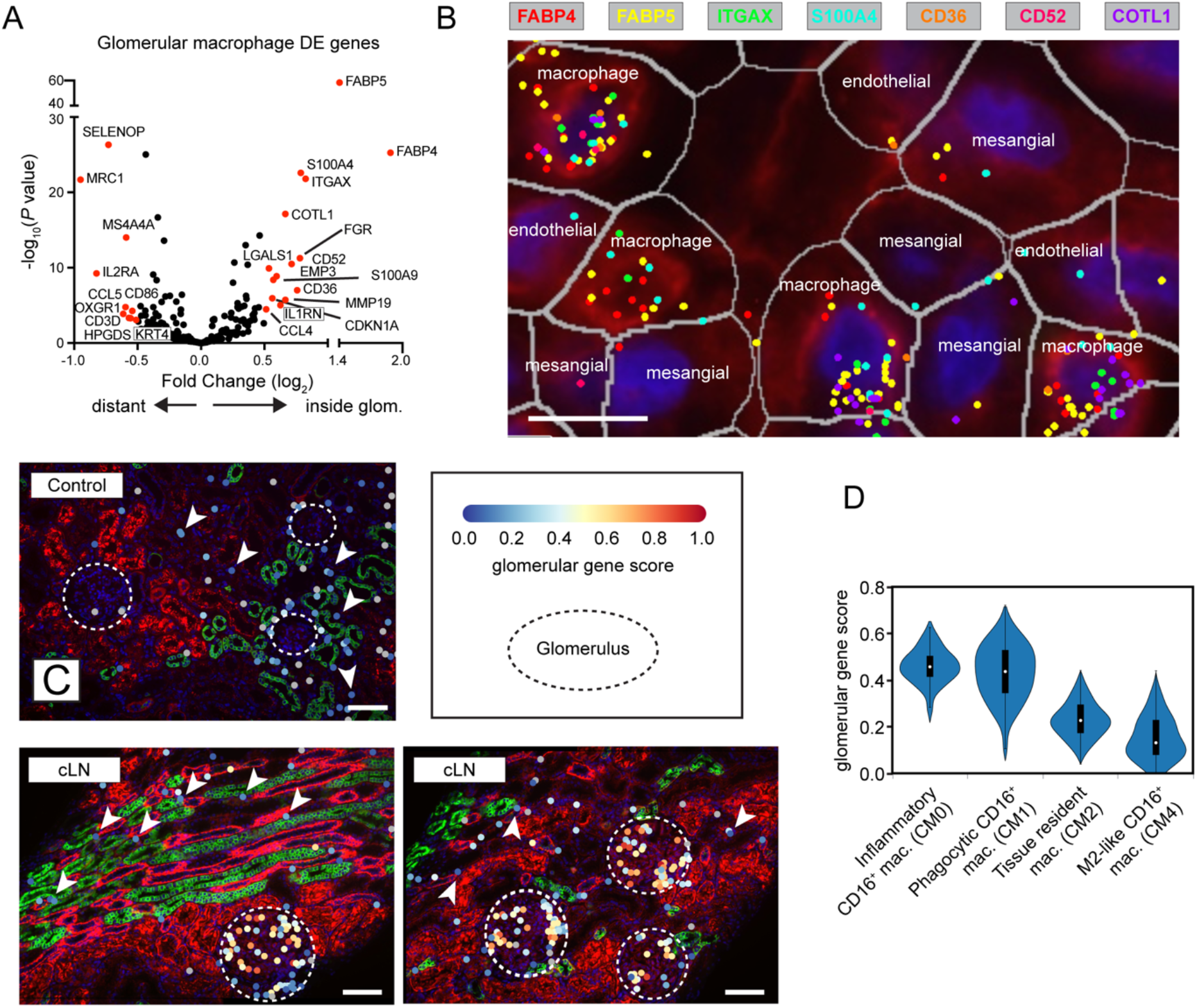
Spatially resolved gene expression in cLN macrophages. **(A)** Volcano plot showing DEG in cLN macrophages residing inside vs. outside glomeruli. **(B)** Expression of representative “glomerulus specific” genes by macrophages residing adjacent to mesangial cells and glomerular endothelial cells in cLN. Scale bars, 10μm. **(C)** Representative images showing increased “glomerular gene score” expression by macrophages residing within cLN glomeruli (dashed circle) vs. tubulointerstitial macrophages (arrowheads). Each dot represents a macrophage colored by metagene expression. Upper panel: representative control sample (Control1). Lower panels: representative cLN (SLE3). Scale bars, 100μm. Image generated using BioTuring Lens. **(D)** Reanalysis of AMP LN dataset (*5*) showing “glomerular gene score” expression by myeloid subpopulations in adult LN.

In a recent study, Arazi et al. distinguished various myeloid subpopulations in lupus nephritis kidneys, with Cluster CM2 most likely corresponding to resident macrophages in a healthy kidney, while Clusters CM0, CM1, and CM4 represented infiltrating monocytes/macrophages (*5*). Further, they detected a putative developmental trajectory from an inflammatory blood monocyte (CM0) to a phagocytic (CM1) and then an alternatively activated (CM4) phenotype. To explore where intra-glomerular macrophages resided along this myeloid trajectory, we devised a “glomerular gene score” based on the top 15 genes upregulated in intra-glomerular vs. extra-glomerular macrophages (*27*). We used BioTuring Lens to visually confirm that the expression of these genes was upregulated in cLN macrophages located within glomeruli (dashed circle), but not control or cLN residing in the tubulointerstitium (arrowheads) (**Fig. 3C**). Notably, reanalysis of the published AMP dataset (*5*) revelated elevated “glomerular gene scores” in both inflammatory (CM0) and phagocytic (CM1) macrophage populations (**Fig. 3D**). Thus, we show using an orthogonal approach that myeloid subsets defined based on scRNA-seq of dissociated tissue are enriched within specific kidney tissue compartments. Whether our data indicate that intra-glomerular macrophages undergo in situ differentiation into CM0 and CM1 or whether distinct functional subsets are present in different patients or individual glomeruli remains to be determined. Nonetheless, our findings highlight how macrophages localized within inflamed cLN glomeruli exhibit distinct transcriptional signatures, implying location-specific roles in cLN disease.

### B cell spatial interaction networks within cLN tubulointerstitial immune hotspots

While current LN histopathological classification is defined by glomerular changes (*10, 28*), LN is also characterized by accumulation of activated B cells, plasma cells, CD4^+^ T cells, plasmacytoid dendritic cells, and myeloid subsets in tubulointerstitial regions (*29*). These cell populations frequently organize into lymphoid-like structures, implying induction of persistent, antigen-driven inflammation within the tubulointerstitium. Ultimately, the failure to attenuate interstitial inflammation results in tubular atrophy and renal fibrosis (*30*). It is likely for this reason that tubulointerstitial inflammation, rather than glomerulonephritis, predicts progression to end-stage renal disease (ESRD) (*31*).

Given the known importance of lymphocytes, and B cells in particular, in LN pathogenesis, we sought to delineate their gene expression patterns and interaction networks. Expanded populations of B lineage cells (B cells and plasma cells) comprised 21% of all immune cells in cLN and were subclustered into 4 groups (B.1 through B.4), defined by differential expression of immunoglobulins, plasma cell markers, and MHC genes (**Fig. 4A, B; Supplemental Table S5**). In contrast to previous scRNA-seq analyses of dissociated kidney tissue (*5*), intermediate gene expression states were detected between B cell subclusters, implying local plasma cell differentiation within the inflamed kidney. Subcluster B.3 cells, which exhibit modest expression of IgG genes and high expression of MHC genes, comprised a larger proportion of B cells in cLN (Chi squared *P=*0.012), whereas B.1 and B.2 cells, which express *IGHM* and *IGHA1* respectively, were more prevalent in controls (Chi squared *P=*0.005 and 0.003; **Fig. 4C**). B cells across all subclusters seldom appeared in isolation, with 95% of them neighboring other immune cells (≥ 1 immune cell within 50 closest neighbors; Fig. 5C) and 38% residing in tubulointerstitial immune hotspots. Within hotspots, B.3 cells were overrepresented, comprising 28% of all B cells and 37% of B cells in hotspots (Chi squared *P*<2.2×10^16^; **Fig. 4D**). Visualization of representative immune foci confirmed spatial interactions between B cells, plasma cells, macrophages, and rare CD4^+^ T cells (**Fig. 4E**). Moreover, plasma cells within local foci expressed IgA, IgG, and IgM immunoglobulin gene transcripts, further supporting local plasma cell generation in the context of an ongoing adaptive immune response in cLN (**Fig. 4F**).

**Figure 4:**
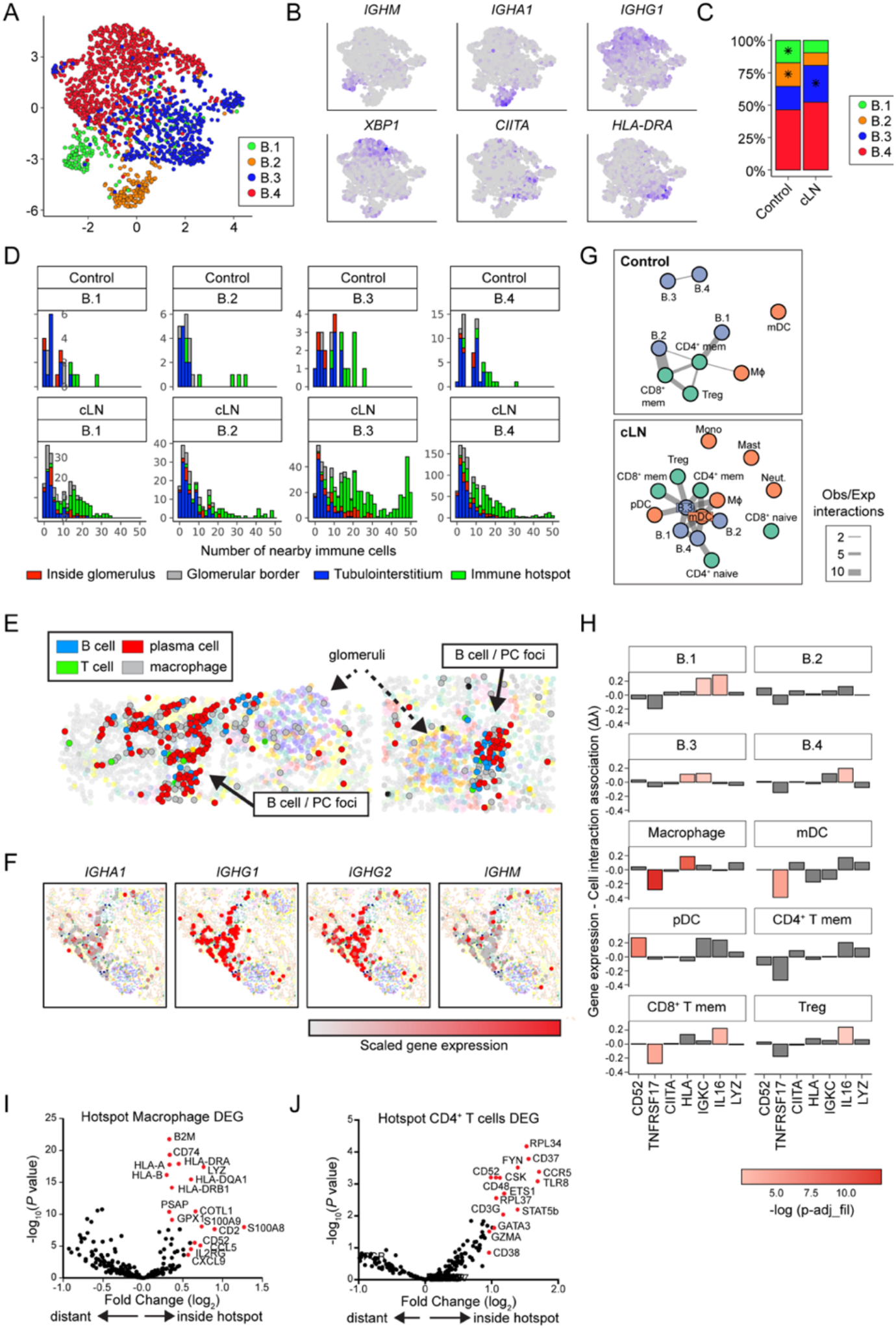
Gene expression and cell interaction patterns within immune hotspots. **(A)** UMAP projection of B cell lineage subclusters B.1 – B.4. **(B)** Heatmap of gene expression in B cell subclusters. **(C)** Distribution of B cell subclusters in control and cLN. \**P*<0.05, by chi-squared test. **(D)** Histograms depicting number of the 50 closest cells to each B cell subset that are immune cells (color coded by B cell location and divided by disease status and B cell subcluster). **(E)** Clusters (solid arrows) of B cells, plasma cells, macrophages, and rare CD4^+^ T cells located with tubulointerstitium of representative cLN kidney. Dashed arrows indicate glomeruli. **(F)** Immunoglobulin gene transcripts (red=scaled gene expression) in cLN plasma cell foci. **(G)** Graphs of interactions between immune cell types that occur more frequently than would be expected by chance, as modelled by a hypergeometric distribution. Cells are colored by lineage and edge thickness is proportional to the ratio of observed:expected number of interactions across all patients. **(H)** Bar chart depicting the relationship between B.3 cell gene (or gene set) expression and the number of interacting cells. Height of the bar is the Poisson regression estimate of that gene (or set of genes). **(I, J)** Volcano plots showing DEG in cLN macrophages (I) and CD4^+^ T cells (J) residing inside vs. outside “immune hotspots”.

**Figure 5:**
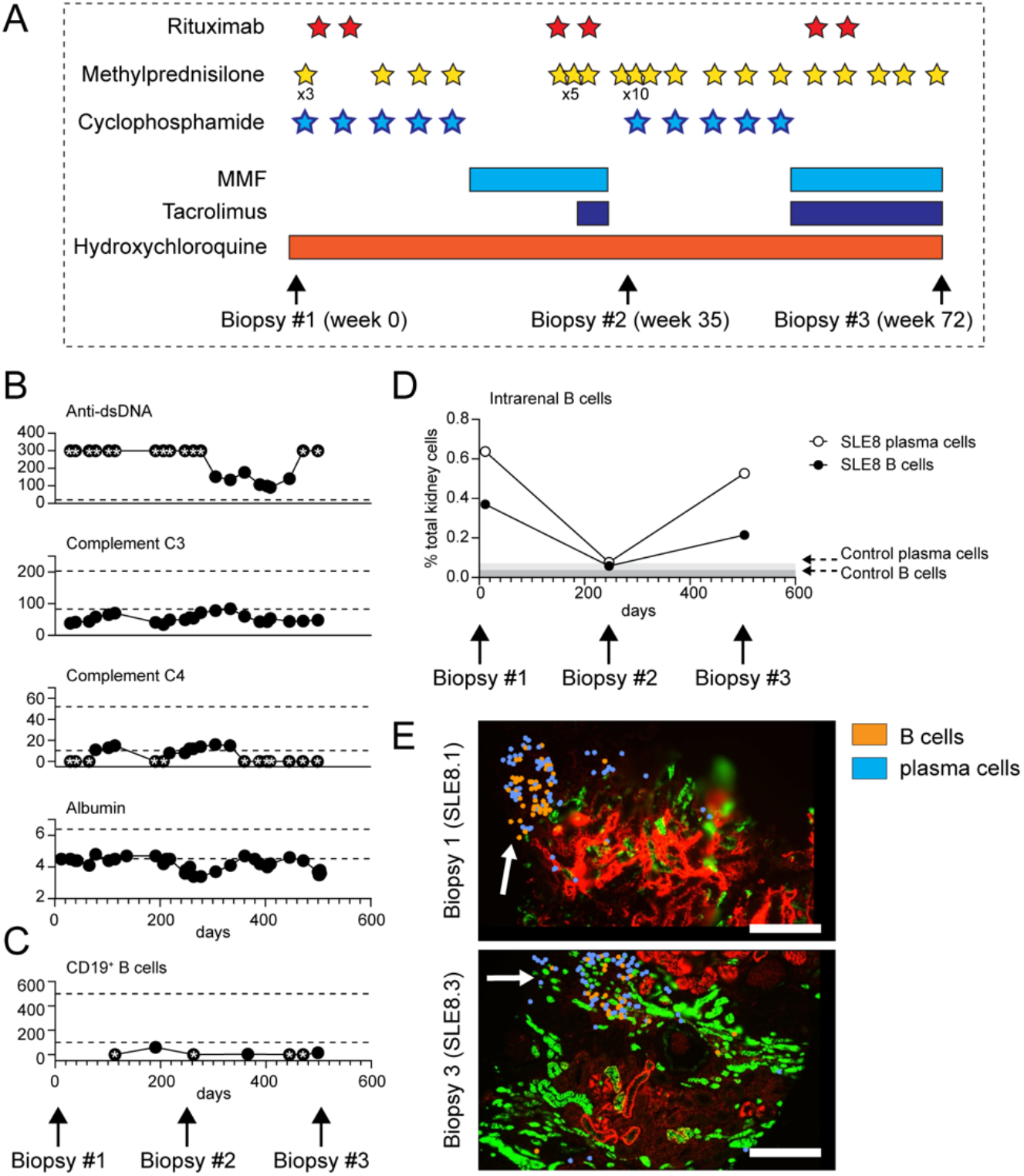
Rituximab-resistant B cell foci in treatment-resistant cLN. **(A)** Diagram depicting immunosuppressive therapies and timing of serial kidney biopsies in subject SLE8. Stars indicate individual doses of rituximab (1000mg, red), IV methylprednisolone (500mg, yellow), and cyclophosphamide (900-1200mg, blue). Bars indicate duration of oral medication use. **(B, C)** Trajectory of clinical biomarkers of cLN disease activity **(B)** and circulating CD19^+^ B cell numbers **(C)** in SLE8. Each dot indicates an individual laboratory value. Stars indicate values above/below the limit of detection. Dashed lines indicate normal range for clinical assay. **(D)** Intrarenal B cells (black circle) and plasma cells (open circle), as percentage of total kidney cells, in 3 serial biopsies from subject SLE8. Grey bars indicate % B cells/plasma cells in control kidney. **(E)** Representative image showing persistent tubulointersitial foci of B cells (orange) and plasma cells (blue) in sample SLE8.1 (obtained prior to rituximab treatment) and sample SLE8.3 (post-rituximab in the setting of circulating B cell depletion). Scale bars, 200μm.

To gain an unbiased understanding of B cell interactions with other immune cells, we performed a network analysis, modelling interactions using hypergeometric distribution. Our analysis revealed the presence of a tubulointerstitial network in cLN, with subcluster B.3 occupying a central node and exhibiting interactions with memory T cells, dendritic cells, macrophages, and other B cell subclusters. In contrast, immune networks in control subjects primarily consisted of memory T cells as central nodes, with minimal contributions from B cell lineages (**Fig. 4G**).

To examine which genes underlie subcluster B.3 interactions in cLN, we used scaled gene expression values to model the number of interactions between B.3 cells and other immune cells, focusing on genes highly expressed in B.3 cells (**Fig. 4H**). B.3 cell expression of MHC genes was associated with macrophage interactions, and the lymphocyte marker *CD52* with pDC interactions. In contrast, B.3 cells with higher expression of *IL16* were more likely to interact with CD8^+^ memory T cells, regulatory T cells, and other B cells (**Fig. 4H**), a notable finding given recent reports identifying urine IL-16 as a biomarker of LN disease activity (*32, 33*). Together with our spatial network analyses (**Fig. 4G**), these data suggest that B cells may both orchestrate and restrain tubulointerstitial immune foci, via IL-16 dependent recruitment of effector T cells (*34*) and by induction of regulatory T cells (*35*).

While B cells/plasma cells were found to predominantly localize within immune hotspots, macrophages and CD4^+^ T cells exhibited more diverse distribution across different anatomical regions of the kidney (**Fig. 2E**). These spatial data facilitated the quantification of differentially expressed genes (DEGs) based on their location within immune hotspots. Interestingly, macrophages residing within these immune foci expressed Class I and II HLA genes, as well as MHC class I component beta-2-microglobulin (*B2M*) and MHC class II chaperone invariant chain (*CD74*), suggesting they play a role in antigen presentation (**Fig. 4I**). Similarly, CD4^+^ T cells within immune hotspots demonstrated upregulation of *TLR8* (linked to LN pathogenesis and expressed by regulatory T cells (*36, 37*)), the Src family kinases (*FYN*, *CSK*), CD52 (expressed by antigen-activated CD4^+^ T cells with suppressive activity (*38*)), and the chemokine receptor (*CCR5*) (**Fig. 4J**). Collectively, these data suggest that spatial interactions between B cells, myeloid cells, and T cells within tubulointerstitial “immune hotspots” exhibit transcript signatures consistent with both adaptive immune functions and induction of regulatory mechanisms.

### Persistent B cell tubulointerstitial foci despite therapeutic B cell depletion in treatment-resistant cLN

Despite the anticipated pathogenic role of B cells in lupus nephritis, the clinical trials evaluating the efficacy of rituximab, a B cell-targeting therapy directed against the CD20 antigen, have yielded disappointing results (*39, 40*). One plausible explanation for this therapeutic failure could be the insufficient depletion of B cells within the inflammatory foci in the kidney (*41–43*). To investigate this hypothesis, we conducted a comprehensive analysis of intrarenal B cells using serial biopsy samples obtained from a subject with treatment-resistant lupus nephritis (SLE8.1 - 8.3). At 13 years old, subject SLE8 presented to an outside institution with Class II LN, for which she was treated with prolonged corticosteroids and rituximab. Her course was complicated by extensive steroid-induced avascular necrosis, but she achieved remission before presenting to our institution with a cLN flare at 16 years old. Repeat diagnostic kidney biopsy (sample SLE8.1) showed Class IV proliferative LN (60% endocapillary proliferation; 5% segmental crescents; no tubulointerstitial fibrosis). Immunosuppression with cyclophosphamide (5 monthly doses) and rituximab was initiated, before transition to oral MMF. Despite aggressive immunosuppression, she developed recurrent proteinuria and active urine sediment which prompted initiation of oral tacrolimus and rituximab re-administration, followed by repeat kidney biopsy (SLE8.2; active Class IV LN with 95% endocapillary proliferation and 40% cellular crescents). Despite a second course of monthly cyclophosphamide and repeat rituximab, remission was not achieved and an additional biopsy (SLE8.3) was obtained (treatment course and timing of biopsies summarized in **Fig. 5A**).

Despite this aggressive immunosuppression, subject SLE8 exhibited persistent lupus disease activity, as evidenced by hypoalbuminemia, hypocomplementemia, and elevated dsDNA antibodies (**Fig. 5B**). Notably, rituximab effectively depleted B cells in the peripheral blood with CD19^+^ B cell counts remaining below the limit of detection in 4 of 7 samples collected over a 16 month period (**Fig. 5C**). However, CosMx spatial transcriptomics of three serial kidney biopsies revealed persistent B cell foci within tissue sections obtained 72 weeks apart (SLE8.1 and SLE8.3), demonstrating the lack of concordance between the peripheral blood and intrarenal B cell numbers (**Fig. 5D, E**). Although limited to an individual patient, these findings provide compelling support for the hypothesis that inadequate depletion of effector B cells within the inflamed renal tissues can account for the lack of rituximab treatment response in lupus nephritis.

### Disease-associated changes in resident glomerular cells in cLN

A significant advantage of CosMx single cell resolution spatial transcriptomics is the ability to perform integrated analysis of infiltrating immune cells and stromal kidney cells within the same tissue section. The glomerular deposition of autoantibody:autoantigen immune complexes promotes both the recruitment of inflammatory cells and the activation of resident glomerular cell subsets. For example, mesangial cells respond to accumulating immune complexes by proliferating and secreting pro-inflammatory chemokines and cytokines, while podocyte injury and loss results in protein leakage and ultimately glomerulosclerosis (*22*). Consistent with this model, the ratio of mesangial cells to podocytes was increased in cLN, indicating mesangial cell proliferation and a relative decrease in podocyte numbers in SLE (**Fig. 6A, B**).

**Figure 6:**
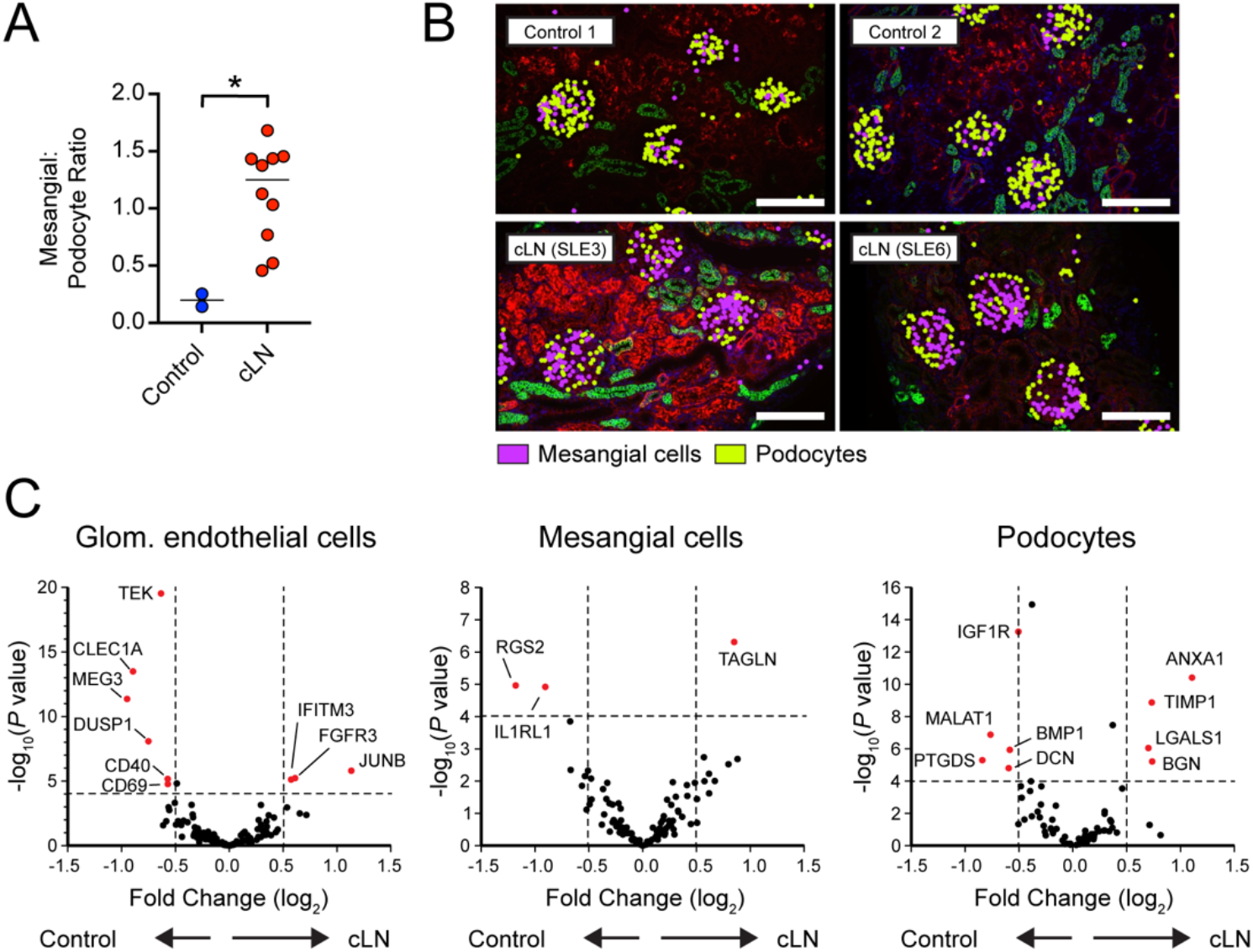
Lupus nephritis is characterized by altered gene expression in resident glomerular cells. **(A)** Mesangial:podocyte ratio in control vs. cLN. * *P*<0.05, by two-tailed Mann-Whitney test. **(B)** Representative images showing mesangial cells (purple) and podocytes (yellow) in control (Control1, Control2) and cLN (SLE3, SLE6) kidney tissues. Morphology markers: B2M/CD298, red; PanCK, green. Scale bars, 200μm. **(C)** Volcano plots showing differentially expressed genes (DEG) in glomerular endothelial cells (left), mesangial cells (middle), and podocytes (right).

Quantification of differentially expressed genes (DEG) uncovered broad transcriptional changes in resident glomerular cells in SLE (**Fig. 6C**). Relative to control cells, cLN glomerular endothelial cells downregulate expression of *TEK* (encoding the angiopoietin-1-binding TEK receptor tyrosine kinase), the angiopoietin-1/TEK signal regulator *DUSP1* (Dual Specificity Phosphatase 1), and *CLEC1A* (encoding the C-type lectin domain family 1 member A), suggesting altered glomerular endothelial function and cross-talk with surrounding mesangial matrix. Among the upregulated genes in cLN mesangial cells, *TAGLN*, encoding transgrelin, is a marker for proliferating mesangial cells responding to tissue injury (*44*). Mesangial cells in cLN also upregulated genes encoding collagen molecules (*COL3A1, COL1A2, COL4A1, COL6A*), matrix metalloproteinases and inhibitors (*MMP14, MMP19, TIMP1*), and chemokines (*CSF1*, *CXCL9*). Conversely, cLN was characterized by downregulation of mesangial cell *RGS2*, a suppressor of renal inflammatory and fibrogenic responses via inhibition of angiotensin II (AngII) and AngII receptor (AT1R) signaling (*45*). In parallel, cLN podocytes upregulated expression of annexin A1 (*ANXA1*), an annexin superfamily member known to promote resolution of inflammation and alleviate kidney injury (*46, 47*) and *LGALS1*, encoding the slit diaphragm component galectin-1 which is increased in diabetic nephropathy (*48, 49*).

### Correlation between glomerular cell gene expression and tissue morphology

The development of LN is not merely variable between affected individuals, but also occurs heterogeneously across kidney tissues sections from individual patients. In the International Society of Nephrology and the Renal Pathology Society (ISN/RPS 2016) classification system (*28*), Class III and Class IV LN are distinguished by the presence of endocapillary hypercellularity involving <50% (Class III) or >50% (Class IV) of sampled glomeruli. Thus, by definition, LN histopathologic features are not uniform across individual tissue sections. We leveraged this heterogeneity to assess whether differential gene expression occurs as a function of disease, LN Class, specific patient, or at the level of individual glomeruli. To do this, we first scored individual glomeruli based on the presence of LN Class-defining lesions and annotated each glomerulus as: 1) “normal” (i.e. lacking any histologic changes); 2) “endocapillary” (i.e. presence of Class IV-defining lesions endocapillary hypercellularity, wire-loops, or double contours); and 3) “other changes” (including isolated mesangial hypercellularity). Next, the average gene expression profile within each glomerulus was taken separately for podocytes, mesangial cells, and endothelial cells. These 3 expression profiles were concatenated, attaining for each glomerulus a profile of cell type-specific expression (**Fig. 7A**). This approach allowed the interrogation of glomerulus-specific gene expression unconfounded by changes in relative cell type abundance. By integrating histologic scores and gene expression profiles from individual glomeruli, we tested whether cLN-specific transcriptional changes are broadly shared across tissue sections or whether altered gene expression occurs at the level of individual glomeruli manifesting lupus-associated pathology.

**Figure 7:**
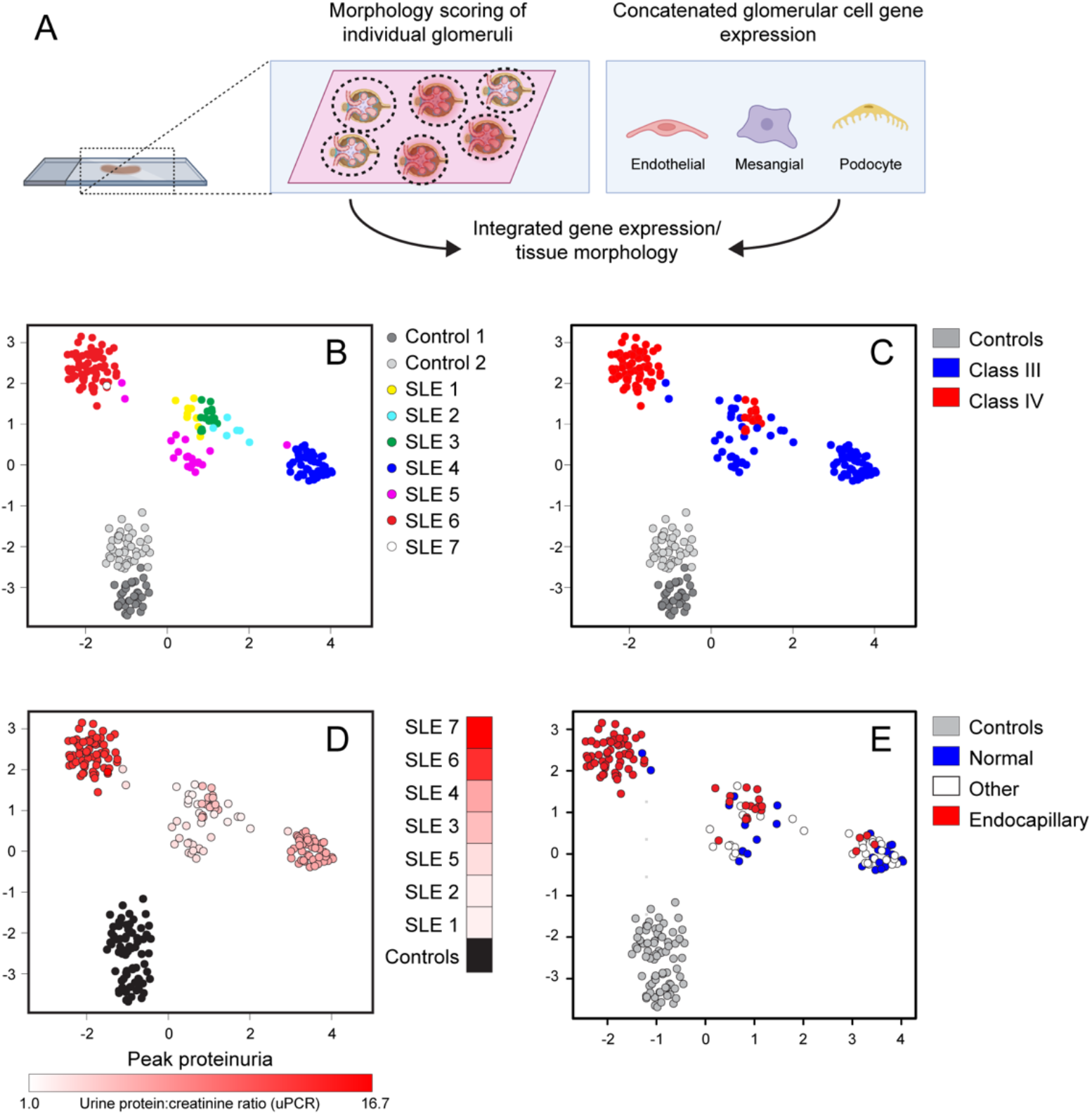
Correlation between glomerular gene expression and tissue histopathology. **(A)** Concatenated glomerular cell gene expression was analyzed based on morphologic scoring of individual glomeruli. **(B-E)** UMAP projections of individual glomeruli colored by patient ID (B), LN Class (C), pre-treatment urine protein:creatinine ratio (uPCR) (D), and lupus-defining histopathology (E). Each dot indicates an individual glomerulus.

By deriving a Uniform Manifold Approximation and Projection (UMAP) of glomerulus-specific gene expression profiles, we demonstrated that glomeruli from individual cLN subjects predominantly clustered together separate from healthy controls (**Fig. 7B**). While Class III and Class IV LN partially segregated into two groups, these categories overlapped (**Fig. 7C**). In addition, we observed a trend towards increased baseline proteinuria in the glomerular cluster derived from SLE6 and SLE7 (**Fig. 7D**). More importantly, the three histology scoring groups were distributed across principal component analysis (PCA) space, with “normal” glomeruli (Group 1) co-localized with “endocapillary” glomeruli (Group 2) from the same patient, rather than with histopathologically similar glomeruli from other subjects (**Fig. 7E**). Indeed, whereas glomerular cell gene expression differed markedly between cLN and controls (**Fig. 6C**), no DEG reached statistical significance when Group 1 (“No path”; normal) and Groups 2 and 3 (“Path”; endocapillary and other) glomerular endothelial cells, mesangial cells, and podocytes in cLN were directly compared (**Supplemental Figure 2**). Thus, alterations in glomerular gene expression are observed across cLN tissue samples, even in morphologically normal glomeruli, indicating a surprising disparity between transcriptional and morphometric characteristics of human cLN.

### Identification of spatially-correlated gene modules linked with induction of inflammation and tissue fibrosis in cLN

To compliment the above hypothesis-driven analyses, we conducted a data-driven unbiased exploration of spatially resolved gene expression in cLN. Using the R package InSituCor (*50*), we identified modules of genes exhibiting spatial correlation, that is the propensity for co-expression in specific anatomical regions beyond what could be accounted for by the cell type distribution alone. This analysis identified 51 distinct modules, each comprised of 3 to 82 genes (**Fig. 8A**). Several interferon-stimulated genes (*ISG*), including *BST2*, *CD81*, *HLA-E*, *IFITM1*, *IFITM3*, *MX1*, and *STAT1*, formed one module (**Fig. 8B**). This observation was striking since this module was identified in an unbiased manner based on spatially-correlated gene expression, but is consistent with known links between type 1 interferon activity and lupus pathogenesis and the ongoing clinical development of type 1 IFN blocking therapies in human (*51, 52*). “ISG module” activity was focused within glomerular cell subsets but varied greatly between controls and SLE, among individual patients, and spatially within samples (**Fig. 8C, Supplemental Figure 3**).

**Figure 8:**
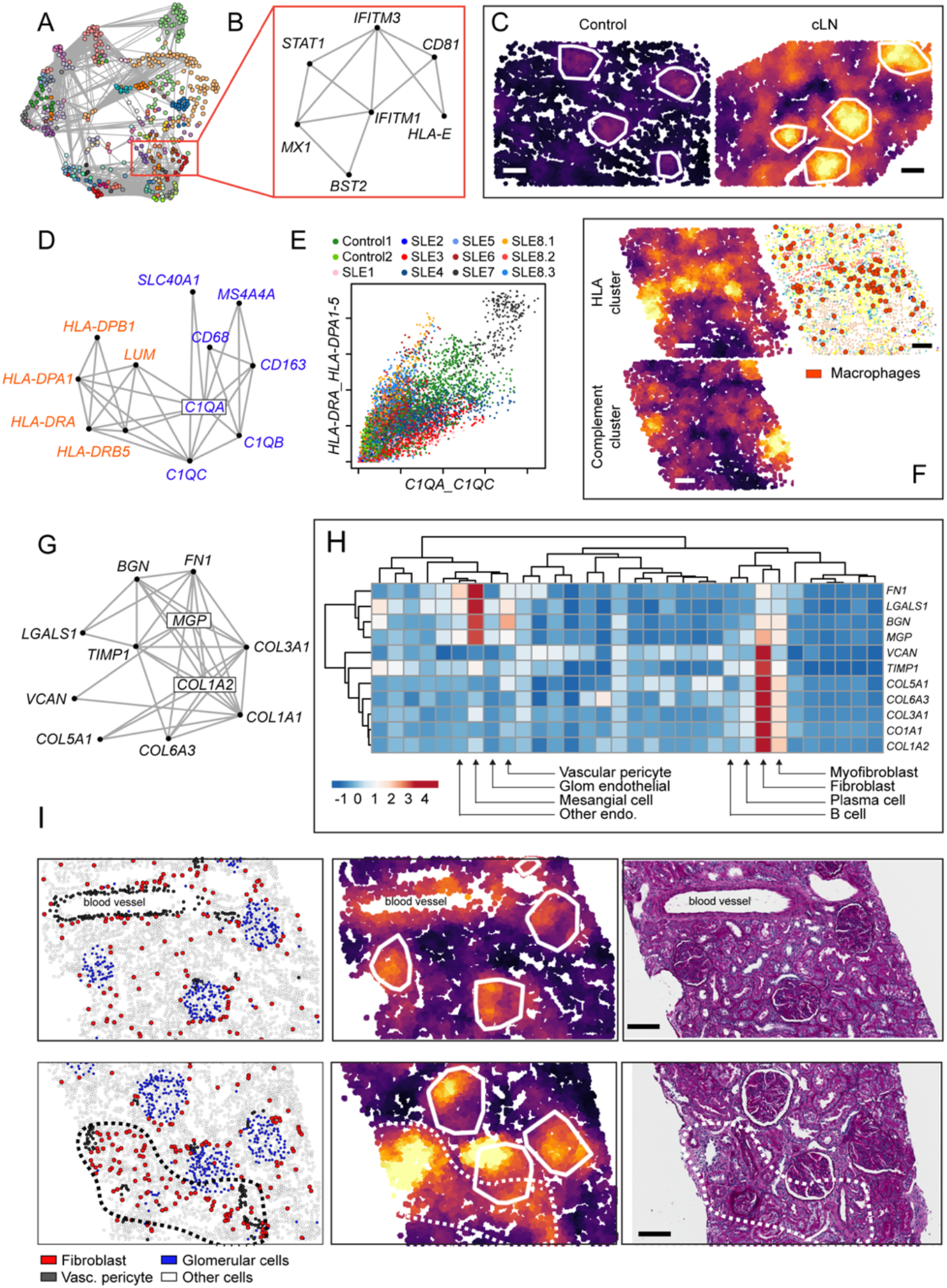
Spatially correlated gene modules in cLN. **(A)** Network connecting spatially correlated gene pairs. Node color shows clusters of mutually correlated genes. **(B)** Spatial correlation network for interferon stimulated gene (ISG) module. **(C)** ISG module activity in representative kidney tissue from healthy control (left) and cLN (right). Scale bars, 100μm. **(D)** Spatial correlation network for two macrophage-associated modules (“Class II HLA cluster”, red; “Compliment cluster”, blue). **(E)** Neighborhood-level activity of each macrophage module across all analyzed cells (colored by sample ID). **(F)** Heatmap showing corresponding “HLA cluster” (upper left), “Compliment cluster” (lower left), and cell type annotation image highlighting macrophage location (red dots) from representative cLN subject. Scale bars, 100μm. **(G)** Spatial correlation network for fibrosis module. (H) Heatmap showing expression of individual fibrosis module genes by kidney cell type. **(I)** Representative cLN tissues sections (SLE6) showing corresponding cell type annotations (left), fibrosis module expression (middle), and Periodic acid–Schiff (PAS)-stained tissue sections (right). Upper panels show fibrosis module expression overlapping with glomeruli and arteriole surrounded by vascular pericytes. Lower panels highlight tubulointerstitial fibrosis module expression within an anatomical region enriched for fibroblasts (dashed line). Scale bars, 100μm.

Spatial domains enriched for macrophages expressed two separate modules with overlapping but distinct tissue distributions. These include one module expressing Class II HLA molecules *HLA-DPA1*, *HLA-DPB1*, *HLA-DRA*, and *HLA-DRB5*, and the extracellular matrix protein Lumican (*LUM*), and another comprising macrophage marker CD68, complement components (*C1QA*, *C1QB*, *C1QC*), tetraspan molecule *MS4A4A*, hemoglobin-haptoglobin scavenger receptor *CD163*, and the iron transporter Ferroportin-1 (*SLC40A1*) (**Fig. 8D-F**). Most tissues were polarized towards one of these clusters, although SLE7 exhibited high-level expression of both modules, driven by cells localized within a dense immune hotspot. Interestingly, the first two samples from the longitudinal subject with treatment-refractory disease (SLE8.1 and SLE8.2) were polarized towards the “Class II HLA cluster”, while in the final sample (SLE8.3) the “Complement cluster” dominated, findings consistent with changes in gene expression within macrophage-rich regions over time and in response to treatment (**Supplemental Figures 4, 5**).

Finally, we observed a gene module potentially linked with induction of tubulointerstitial fibrosis. This module included collagen molecules (*COL1A1*, *COL1A2*, *COL3A1*, *COL5A1*, *COL6A3*), and matrix components (fibronectin-1 (*FN1*), galacetin-1 (*LGALS1*), biglycan (*BGN*), matrix Gla protein (*MGP*), TIMP metallopeptidase inhibitor 1 (*TIMP1*), and versican (*VCAN*)) (**Fig. 8G**). Interesting, analysis of underlying cell distributions indicated the presence of two submodules: one focusing on mesangial cells and vascular pericytes (*FN1, LGALS1, BGN,* and *MGP*); and a second on fibroblasts, myofibroblasts, and to a lesser extent B cells and plasma cells (*VCAN, TIMP1, COL1A1, COL1A2, COL3A1, COL5A1,* and *COL6A3*) (**Fig. 8H**). Comparing the distribution of kidney cell types and “fibrosis module” expression in a representative cLN sample confirmed these genes were upregulated in kidney regions comprising mesangial cells (glomeruli), around blood vessels (vascular pericytes), and in tubulointerstitial regions enriched with fibroblasts (**Fig. 8I; Supplemental Figure 6**). Notably, “fibrosis module” expression was observed in kidney regions lacking overt tubulointerstitial fibrosis on histopathology. In addition, upregulation of this gene module was also observed in a control samples (Control1; nephrectomy for chronic vascular congestion) (**Supplemental Figure 6**), findings underscoring that even kidney tissue deemed “histologically normal” on light microscopy can exhibit changes in gene expression identifiable through spatial transcriptomics.

“Fibrosis module” expression patterns also varied between individual cLN samples (**Supplemental Figure 6**). In certain tissues, module expression was concentrated in hotspots smaller than individual glomeruli, corresponding with regions of mesangial expansion, while in other samples (SLE7, SLE8.3) module expression spanned the width of the FOV (0.76mm). In addition, several cLN subjects (SLE6, SLE8.2) exhibited module intensity that peaked within glomeruli, while in others (SLE4, SLE5, SLE7) expression was focused in the tubulointerstitium. Since our cLN cohort focused on newly diagnosed disease, these data indicate that “fibrosis-related” gene upregulation can occur early in the cLN disease course and before tubulointerstitial fibrosis is evident by light microscopy. While the clinical significance of “fibrosis module” expression remains to be determined, we hypothesize that these data might explain the rapid increase in histopathologic damage indices noted in repeat kidney biopsies obtained after LN induction immunosuppression (*53*).

## Discussion

Leveraging an advance in single cell spatial transcriptomic technology we profiled the cellular and molecular landscape of cLN and control kidney tissue. These analyses yielded several important insights into the biology and spatial heterogeneity of kidney inflammation in SLE. First, in additional to confirming an anticipated increase in infiltrating immune cells in cLN (*5*), we show that immune cell transcriptional profiles vary as a function of tissue distribution, such as expression of genes linked with induction of inflammation, phagocytosis, and antigen presentation by myeloid cells located within cLN glomeruli or tubulointerstitial immune hotspots, respectively. Second, we measured gene expression by resident kidney cell types down to the level of individual glomeruli. These analyses uncovered a surprising disconnect between histopathologic changes and local transcriptional profiles. Despite SLE being a systemic autoimmune disease, a surprising feature of LN pathology is that morphologically normal glomeruli can reside adjacent to inflamed glomeruli within the same tissue section; a phenomenon central to the ISN/RPS 2016 LN classification system (*28*). When comparing spatially-resolved gene expression with corresponding histology scores, we observed greater variation in glomerular cell DEG between subjects rather than between glomeruli on the same section, such that even “normal” cLN glomeruli exhibit marked changes compared with controls.

Finally, using a network analysis of cell:cell interactions, we show that tubulointerstitial B cells form a central node within tubulointerstitial immune hotspots in cLN. The proximate location of B cells, plasma cells, memory CD4^+^ T cells, and myeloid lineages within immune hotspots lends support to a model in which intrarenal B cells undergo local expansion and differentiation into plasma cells. While confirmation of this hypothesis will require additional data, such as measurement of clonal relatedness of spatially collocated B cells, these findings support clinical development of B cell targeted agents in LN (*21, 54*). Moreover, our analysis of serial kidney biopsies from an adolescent with treatment-resistant cLN provides mechanistic insights into the limited clinical efficacy of the anti-CD20 monoclonal rituximab in LN (*40*). Whereas circulating B cells are rapidly depleted following CD20 targeting, post-rituximab persistence of B cells within inflamed target organs and secondary lymphoid tissues has been documented in both murine lupus models (*41*) and in humans with diverse humoral autoimmune diseases (*42, 43*). Thus, a likely proximate cause of rituximab resistance in our patient and in subjects enrolled in the LUNAR trial is relative inefficacy of B cell depletion at the site of kidney inflammation. For this reason, agents exhibiting more effective B cell depletion, such as the type II anti-CD20 monoclonal Obinutuzumab (*21*) or CD19-directed chimeric antigen receptor (CAR) T cells (*54*), are undergoing clinical development in LN. Given the feasibility of performing spatial transcriptomics on FFPE tissue, we anticipate that analysis of post-treatment kidney biopsies to confirm anticipated pharmacologic effects will serve as an important adjunct for both patient management and LN clinical trials.

The CosMx platform supported a rich set of queries into lupus biology using relatively few cLN tissue samples. Using typical spatial analysis approaches, we generated cell type maps and described spatial neighborhoods. The ability to profile a large number of genes at single cell resolution was a boon, markedly improving the precision of our maps and uncovering important details such as immune lineage location within distinct anatomical compartments. Although 960 genes represent a fraction of the total transcriptome, this panel was sufficient to annotate of all major kidney cell types and to uncover many highly significant genes in our comparisons. We could also ask questions unanswerable by single-cell or spatial data alone. Because we could simultaneously measure single cells and their surroundings, we could evaluate how cell phenotype interacted with the local environment, a critical advantage of single cell spatial transcriptomics platforms over other single cell and spatial transcriptomic methods.

Since single-cell spatial transcriptomic platforms are relatively new, the field has yet to establish standardized analysis methodologies. We found two contrasting approaches to be productive. First, we performed analyses driven by biological knowledge. We partitioned the kidney tissue into pre-defined anatomical categories informed by renal anatomy and visual examination of cell type maps. Then, much as an epidemiologist would for patient-level data, we fit carefully specified multivariate models for how cells might respond to their spatial contexts. Although easy to interpret, these data only uncovered trends we explicitly modeled and served as an approximation for all sources of spatial variability impacting gene expression. For this reason, we developed a second data-driven analysis strategy. By searching for genes that shared spatial correlations, without dependence on biological priors, we discovered rich and nuanced behavior beyond what we considered in our multivariate models. Given the number of spatially correlated gene modules we identified, interpreting and validating these results will require a combination of larger patient cohorts, domain knowledge, and correlation with long-term clinical outcomes.

Our study has several limitations. First, our cLN cohort was relatively small and focused specifically on Class III and IV disease, thereby limiting conclusions that we could draw regarding the transcriptional phenotype of other LN Classes and additional pathologic entities, such as thrombotic microangiopathy (TMA) and lupus podocytopathy. Second, except for SLE8, all subjects in the cLN cohort responded to standard immunosuppression regimens, as evidenced by resolution of proteinuria by 6 months (**Supplemental Table S4**). Thus, we were unable to identify predictors of poor prognosis, data with increased clinical relevance given expanded options for adjunctive LN treatment after FDA approval of belimumab and voclosporin (*2–4*). We predict that generating spatial transcriptomic datasets using larger cLN cohorts as well as profiling repeat kidney biopsies obtained after immunosuppression will assist in the identification of markers of treatment response and resistance, thereby informing the selection of initial immunosuppression in cLN.

Despite these caveats, our current dataset offers provocative insights into the biology of human LN. We describe parallel immune networks residing within glomeruli and tubulointerstitial immune hotspots in cLN, each expressing genes linked with both pro-inflammatory and counter-regulatory immune functions. In addition, we observed upregulation of fibrosis related genes in a pediatric cohort with recent cLN disease onset. These findings are consistent with the observation that LN kidneys accrue damage early during treatment (*53*), and have important treatment implications, arguing for aggressive initial immunosuppression in cLN and the need to incorporate adjunctive anti-fibrotic therapies. In addition to informing the biology of human LN, we anticipate that this spatially-resolved cell atlas of cLN will become an important resource for lupus investigators in academia and industry.

## Materials and Methods

### Study Design

The aim of this study was to uncover pathophysiologic mechanisms in cLN using single cell resolution spatial transcriptomics. Research protocols were approved by the Seattle Children’s Research Institute (SCRI) Institutional Review Board. Participants or their parents provided written informed consent before participation in the study. Archived clinical biopsy samples were selected from 7 patients with proliferative Class III or Class IV LN as well as a single subject who underwent three serial biopsies for treatment resistant disease. Control kidney tissue was obtained from nephrectomy tissue for non-inflammatory indications (Control1: chronic vascular congestion; Control2: normal tissue adjacent to renal cell carcinoma). All spatial transcriptomic data were analyzed, and no outliers were excluded. No power calculations were performed. All experiments and data analysis were performed by investigators blinded to sample demographics.

### Single cell spatial transcriptomics

Spatial transcriptomic profiling of FFPE kidney sections was performed using the CosMx Spatial Molecular Imager, as previously described (*9*). A development version of the CosMx Human Universal Cell Characterization RNA Panel was used to measure transcript expression, including the following genes:

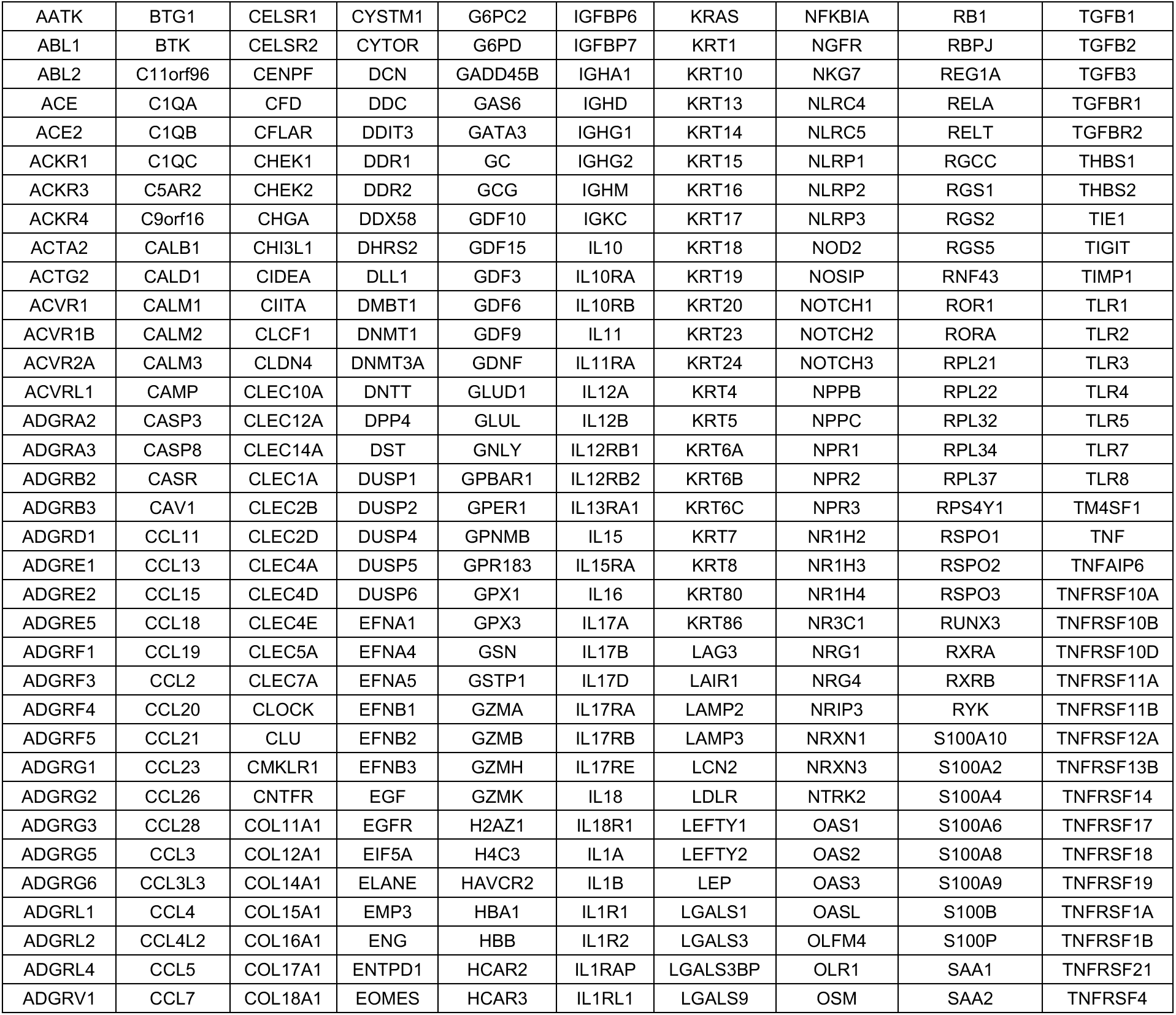
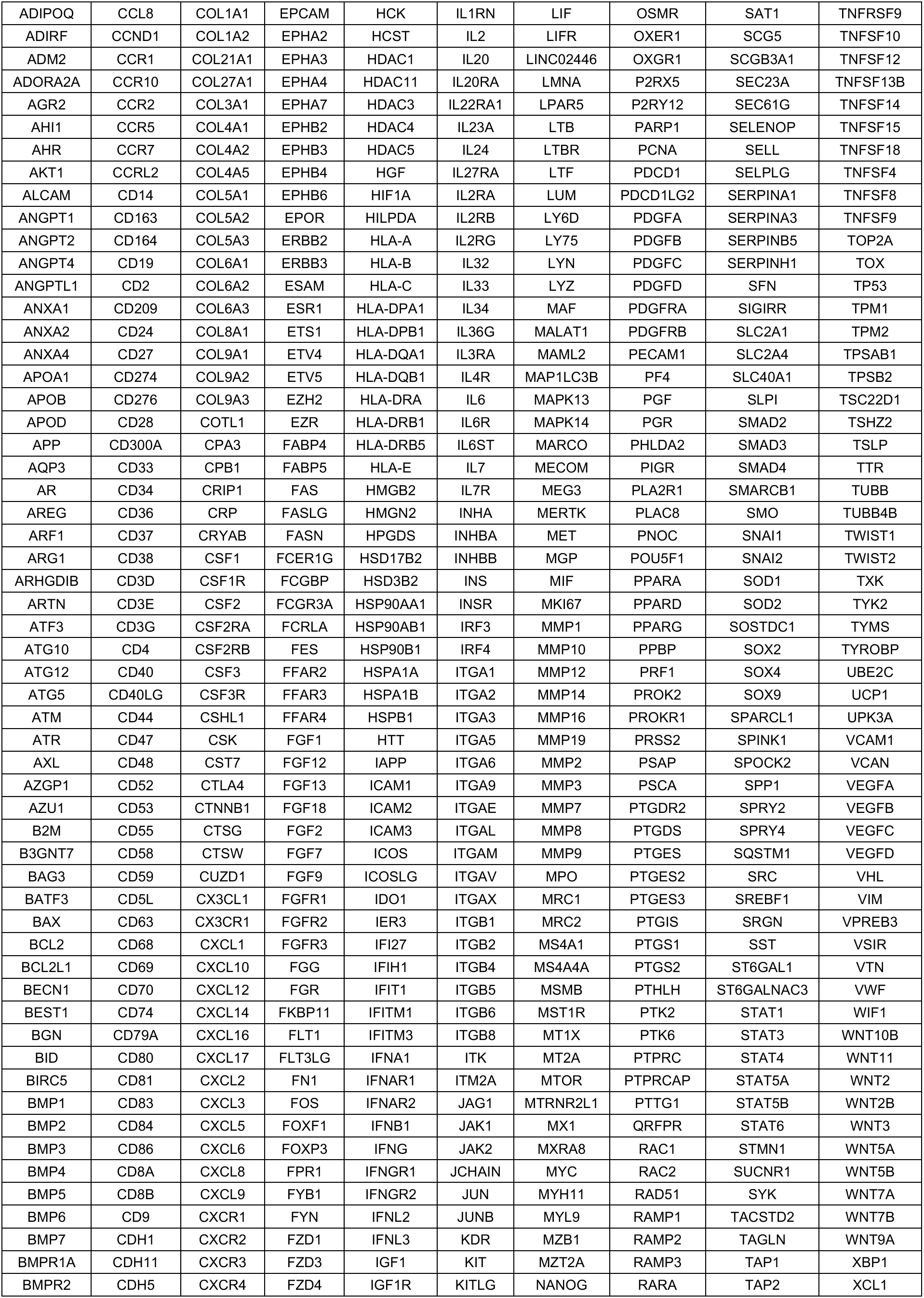
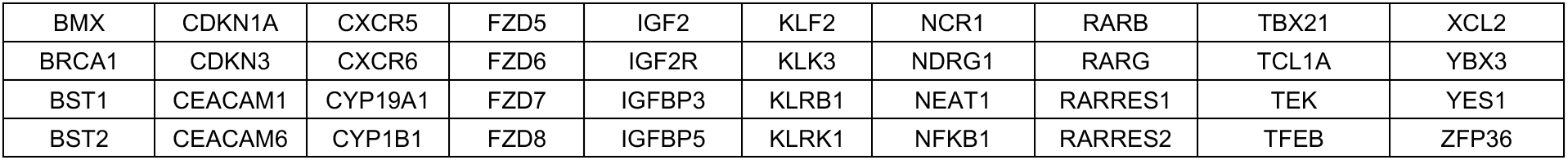

### Cell type classification

A reference matrix of expected cell type expression profiles was constructed based on the Human Cell Atlas kidney dataset, with immune cell profiles replaced by profiles from (*15*). Supervised cell typing was then performed using the InSituType R library (*13*). InSituType was run using gene expression profiles, cell spatial context, and imaging data (size, shape and intensity of immunofluorescence stains for DAPI, CD45, PanCK, CD20, and CD298). Initial cell annotation revealed a “myofibroblast” cell type occupying 3 distinct spatial contexts and regions of the UMAP projection. These were then sub-clustered to generate 3 cell types: “mesangial cells”, “myofibroblasts”, and “vascular pericytes”. The cell type annotated “distinct proximal tubule 1” was similarly sub-clustered into distinct “proximal tubule” and “parietal epithelium” populations based on spatial distribution. All proximal tubule cell types were then combined into a single “PCT” designation.

### Computational definition of glomeruli

Glomeruli were defined through a combination of code-driven detection and refinement by pathologist review. First, the dbscan algorithm was applied to the spatial locations of glomerular cells (podocytes, mesangial cells and glomerular endothelium cells), returning spatially distinct clusters, presumably glomeruli. Only clusters with at least 20 cells were considered. Clusters including multiple neighboring glomeruli by visual inspection were subclustered using the R Mclust package, which naturally fits the ellipsoid shape of glomeruli. The computational glomerulus calls were compared to corresponding Periodic acid–Schiff (PAS)-stained tissue sections by a board-certified pediatric pathologist (R. Reed). Based on this comparison, certain computationally-defined glomeruli were merged and borders between glomeruli were refined. Histopathologic scoring of LN disease activity in individual glomeruli was performed by a board-certified pediatric pathologist (R. Reed), who was blinded to spatial transcriptomic data. Cell position with respect to glomeruli was determined as follows. First, convex polygonal boundaries were fit around each glomerulus using the R function “chull”. Cells inside those polygons were annotated as inside glomeruli. Of the remaining cells, those within 0.05mm of a glomerulus-specific cell type were called as “bordering glomeruli”, or else designated as “tubulointerstitium”. Immune hotspots were defined as groups of at least 50 immune cells with at most 0.05 mm distance from their nearest neighbor. Such groups were detected by clustering immune cell locations using the dbscan algorithm.

### Differential expression analyses

Differential expression in spatial transcriptomics is complicated by imperfections in cell segmentation: if a cell’s border is drawn incorrectly, mRNA transcripts from neighboring cells can be inappropriately assigned to it. Even low-frequency segmentation errors can have pernicious effects on differential expression. For example, T cells in glomeruli will include contaminating signal from neighboring glomerular cells, while T cells surrounded by proximal tubule cells will be contaminated by transcripts from those cells. A comparison of T cells between those spatial contexts would then find spurious significance for glomerular cell and tubule cell marker genes, injecting numerous false positives atop the list of most significant genes.

We employed two countermeasures to combat this effect. First, for each cell type, we only performed differential expression analysis for genes at low risk for contamination from neighbors. We reasoned that a gene with low expression in a cell type and high expression in its neighbors is at greatest risk of contamination. Thus, within each cell type, we compared each gene’s mean expression within the analyzed cell type with mean expression in neighboring cells of separate type, and then filtered out genes with a low ratio.

As a second countermeasure, we included a term in our models that explicitly accounts for contamination. For a given gene, each cell was scored for the total counts of that gene in neighbors of different cell types. (We did not measure expression in neighbors of the same cell type because we did not want to adjust out real biological effects.) This total neighboring expression value was then RankNorm transformed and entered into differential expression models.

Differential expression modeling was then performed with negative binomial mixed effects regression, with raw counts per cell as the outcome. Fixed effects were specified for all variables of interest, and for the neighboring expression term described above. Tissue ID was specified as a random effect. Finally, to normalize out the wide range of signal strength found in different cells, each cell’s log-transformed total counts was included in the model as an offset.

### Differential expression analysis of macrophages

This analysis only considered macrophages in cLN samples. The following variables were included as fixed effects in the model: position vs. glomerulus (inside, bordering, tubulointerstitium, immune hotspot). Tissue ID was included as a random effect.

### Macrophage “glomerular gene score” quantification

The top 15 genes upregulated in intra-glomerular vs. extra-glomerular macrophages were used to calculate a “glomerular gene score”. Using the BioTuring Lens platform, gene set activity in individual macrophages was calculated using the AUCell algorithm (*27*). AUCell measures the activity of the gene set in each cell based on the ranking of all genes, with high AUCell score indicating enriched expression in an individual cell.

### Differential expression analysis of glomerular cells

The following variables were included as fixed effects in the model: disease status (control glomeruli; cLN “No path” (Group 1, normal); and cLN “Path” (combined Groups 2 and 3, endocapillary and other). Tissue ID and glomerulus ID were included as random effects. cLN glomeruli were classified within the “Path” group if the pathologist detected any visible cLN morphologic changes, including segmental sclerosis, mesangial proliferation, endocapillary proliferation, wire loop lesions, circulating immune cells, or other abnormal features. This scheme classified 39 cLN glomeruli without involvement (Group 1) and 226 with visible involvement (combined Groups 2 and 3).

### Spatial correlation analysis

Spatial correlation analysis was performed with a development version of the R package InSituCor (*50*). InSituCor was run on normalized count data, adjusted for the effects of cell type, tissue ID and single cell counts, to uncover spatial correlations beyond those accounted for by these variables.

### B cell subclustering and marker gene analysis

B cells and plasma cells, as designated by the cell type classification algorithm described above, were subclustered using the InSituType algorithm (*13*). The following were included as additional variables used to subcluster cells: cell area, cell aspect ratio, mean CD298 staining intensity, mean PanCK staining intensity, mean CD45 staining intensity, mean CD20 staining intensity, and mean DAPI staining intensity. The Seurat package v4.0.6 (*55*) was subsequently used to identify marker genes for each subcluster (B.1 through B.4). Only genes at low risk for contamination (gene contamination score cutoff ≥ 0.3 for either B cells or plasma cells) were subject to subclustering and marker gene analyses.

### Immune cell-cell interaction analysis

Interacting cells were defined as those within 0.1mm. We sought to assess whether *k*, or the observed number of interactions between any two cell types *A* and *B* with subpopulation sizes *n*_*A*_ and *n_B_*, occurred more frequently than would be expected by chance in a total population of cells of size *n_t_*. Expected interaction frequencies were modelled using the expected median value of a hypergeometric distribution defined by *n*, or the observed number of interactions between all cells, and two other parameters:

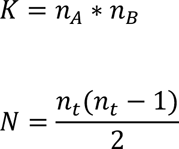

Where *K* is the number of theoretically possible interactions between cell types *A* and *B* interactions, and *N* is the number of theoretically possible interactions between all cells of all cell types. Notably, if cell type *A* = *B* (i.e. one models the expected interaction frequency between cells of the same type), then:

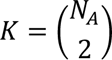

Finally, the expected hypergeometric distribution was used to compute one-tailed p-values for *k*, or the observed number of interactions between cell types *A* and *B*.

Because cells in different patients have no chance of interacting with one another, values of *N*, *K*, *n*, and *k* were computed for each patient individually and then summed across all patients prior to performing the analysis described above.

### Modelling the relationship between B.3 gene expression and B.3 cell-immune cell interaction frequency

Only genes specifically upregulated in B.3 cells and B.3 interaction partners that interact more frequently than would be expected by chance were considered in this analysis. To avoid multicollinearity, genes were first clustered into 7 groups of coregulated gene sets. Gene expression values within each gene set were used to create gene set scores for each cell, which were themselves subjected to Z-score scaling to allow for direct comparison of regression coefficient estimates. Outlier gene set score values (Z > 3 or Z < -3) were removed. A Poisson regression was used to model the number of interactions between B.3 cells and each other cell type according to the expression of gene scores G1 through G7. Patient identity was included as a covariate in the model to regress out any patient-specific cell interaction effects (e.g. differential sampling of tubulointerstitial hotspots between patients).

## List of Supplementary Materials

Figures S1 to S6.

Tables S1 to S5.

## Acknowledgments

The authors gratefully acknowledge the patients and their families for participating in this study. We thank employees at BioTuring, Inc. for incorporating CosMx data into the BioTuring Lens platform and troubleshooting data interpretation.

## Funding

This work was support by National Institutes of Health grants:

- R01AR073938 (to S.W.J)
- R01AR075813 (to S.W.J)

The content is solely the responsibility of the authors and does not necessarily represent the official views of the NIH.

Additional funding support provided by the:

- Seattle Children’s Research Institute (SCRI) Center for Immunity and Immunotherapies (CIIT) (S.W.J)

## Author contributions

- Conceptualization: PD, NH, KH, CEA, RCR, DMO, SKB, SWJ.
- Methodology: PD, NH, EDN, SWJ.
- Software: PD, NH
- Validation: PD, NH, SWJ.
- Formal Analysis: PD, NH, EDN, SWJ.
- Investigation: PD, RCR, SKB.
- Resources: KH, NR, RCR, SKB, DMO, SWJ.
- Data Curation: PD.
- Writing – original draft: PD, NH, SWJ.
- Writing – review & editing: PD, NH, EDN, KH, NR, CEA, RCR, DMO, SKB, SWJ.
- Visualization: PD, NH, SWJ
- Supervision: PD, SWJ.
- Project administration: PD, SKB, SWJ.
- Funding acquisition: SKB, SWJ
- Supervision: PD, SWJ

## Competing interests

The authors declare the following competing interests.

P.D. is an employee and shareholder of NanoString Technologies, Inc. and has filed a provisional patent covering the algorithm for spatial correlation analyses described in this manuscript. N.H. no disclosures. E.D.N: no disclosures. K.H. received an unrestricted educational grant from Pfizer Global medical grants to support a quality improvement program for patients with juvenile idiopathic arthritis (JIA) which was unrelated to the current study. N.R.: no disclosures. C.E.A.: no disclosures. R.C.R: no disclosures. D.M.O. is a consultant for a Horizon Therapeutics Advisory Board. S.K.B. is an employee and shareholder of Sanofi. S.W.J. is a consultant for Bristol-Myers Squib and previously served as a consultant for Variant Bio and ChemoCentryx, Inc.

## Data and materials availability

After publication, all reasonable requests for materials and data will be fulfilled and all data necessary to understand and evaluate the conclusions of the paper will be archived in an approved database.

---

**Supplemental Figure 1:**
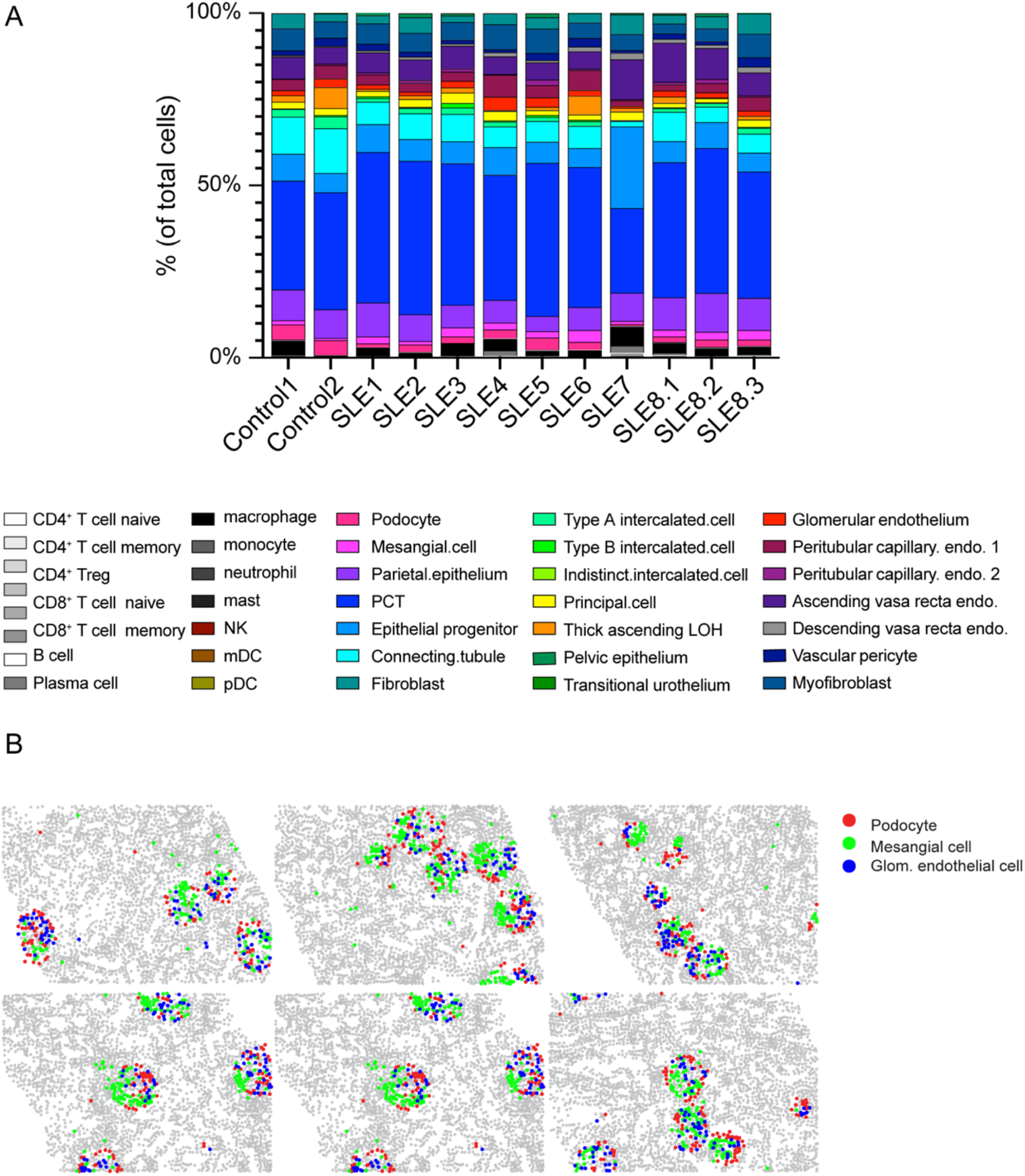
Validation of CosMx cell annotation in human kidney. **(A)** Distribution of 35 kidney and immune cell subsets in healthy control (Control1 / Control2) and childhood-onset lupus nephritis (SLE1-SLE8) kidney tissues. **(B)** Spatial distribution of podocytes (red), mesangial cells (green), and glomerular endothelial cells (blue) in representative cLN kidney tissue. These known glomerular cell types are spatially colocalized with circular structures representing glomeruli. Grey dots: unlabeled cells. Abbreviations: Treg, regulatory T cell; NK, natural killer cell; mDC, myeloid dendritic cell; pDC, plasmacytoid dendritic cell; PCT, proximal convoluted tubule; LOH, loop of Henle; endo, endothelium.

**Supplemental Figure 2:**
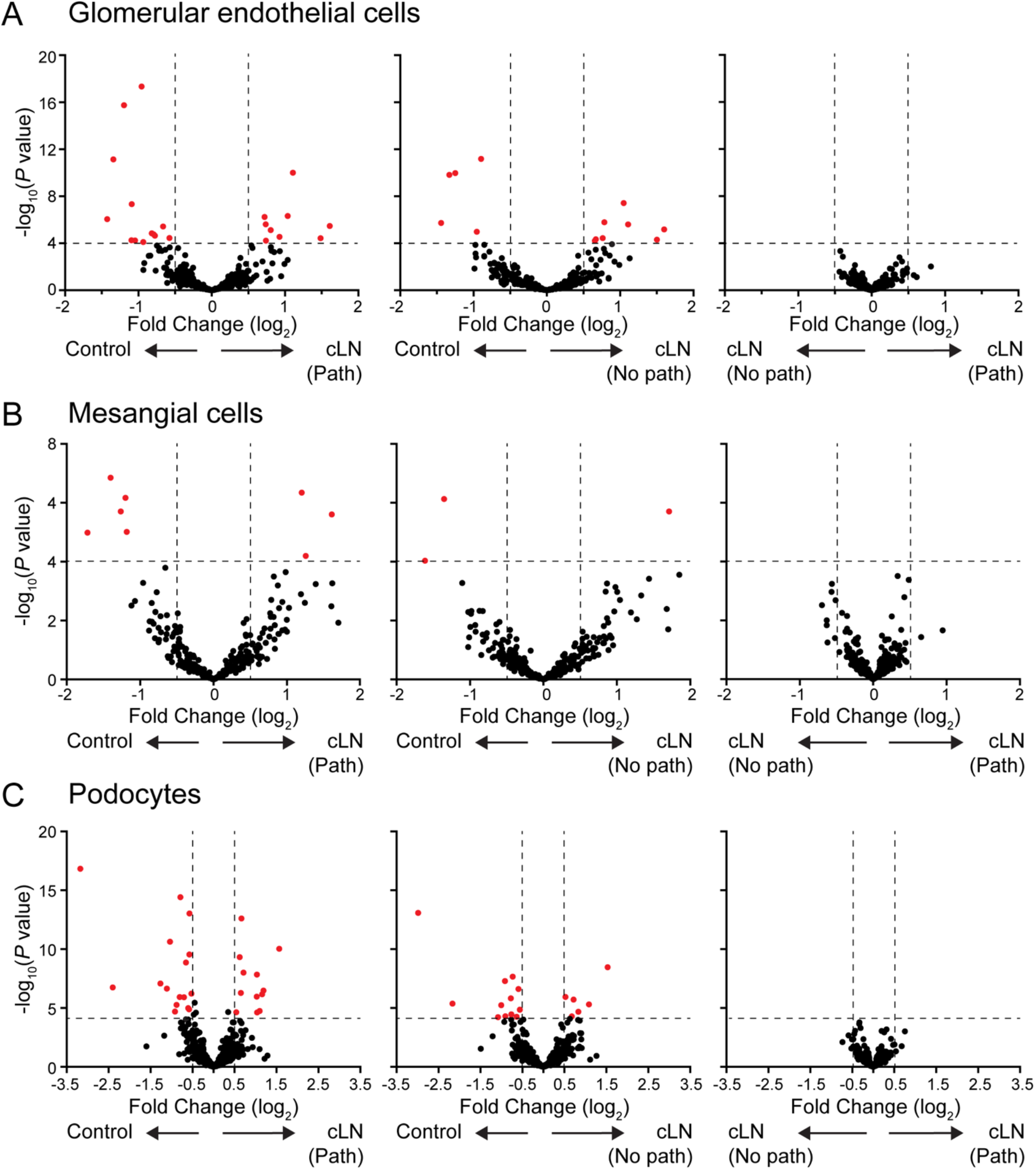
Differential gene expression by glomerular cell subsets based on histologic scoring of individual glomeruli. **(A-C)** Volcano plot showing number of differentially expressed genes (DEG) in glomerular endothelial cells (A), mesangial cells (B), and podocytes (C) as a function of whether cell populations reside within control glomeruli (Control), cLN glomeruli exhibiting no pathologic changes (No path; Group 1), or cLN glomeruli exhibiting any lupus-associated morphologic changes (Path; Groups 2 and 3). Red dots indicate DEG above a threshold of: Log2(fold change) < -0.5 or > 0.5; and -log10(*P*)>4.

**Supplemental Figure 3:**
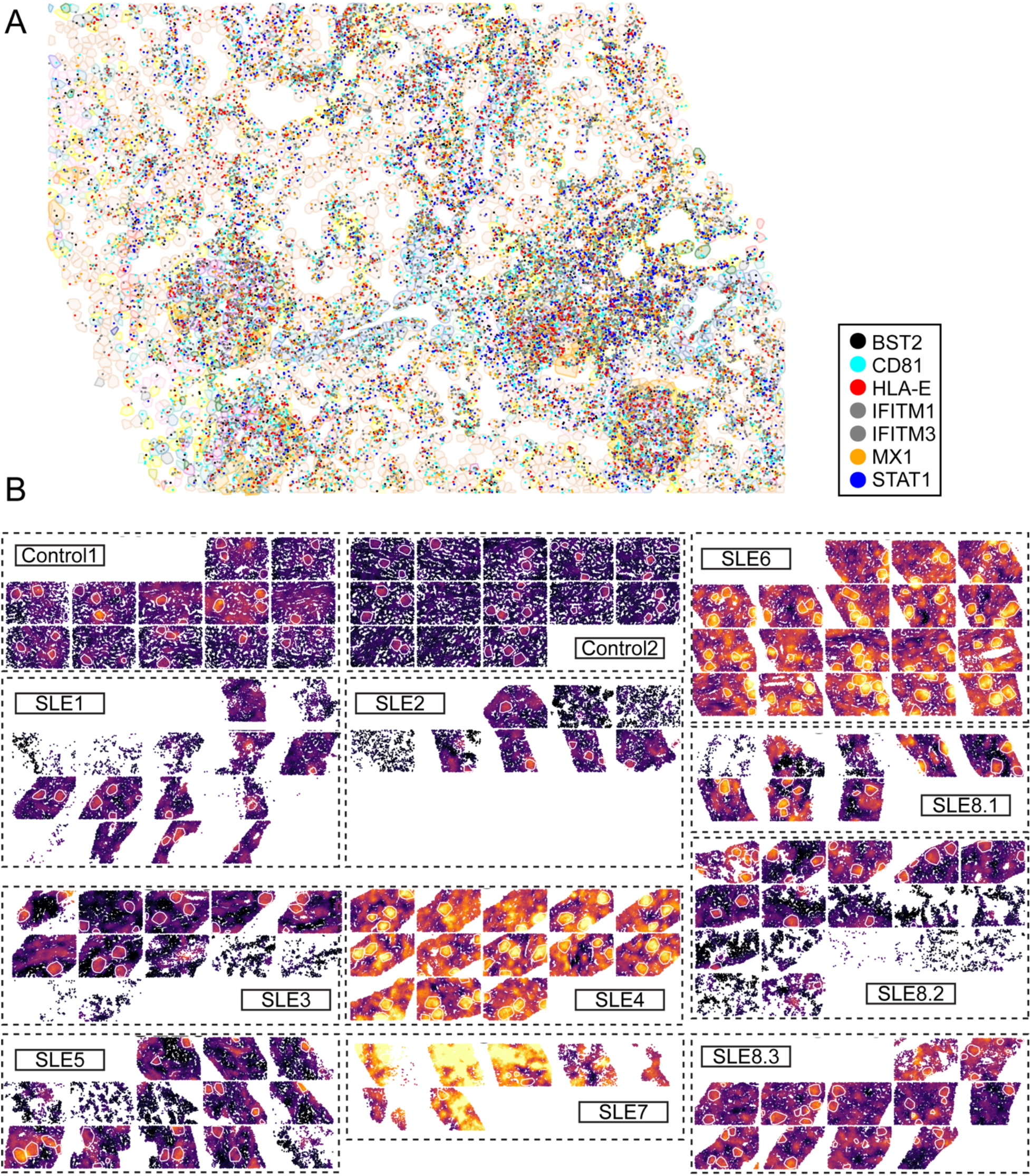
ISG spatial module in cLN. **(A)** mRNA locations of genes within the interferon stimulated gene (ISG) module in representative cLN kidney tissue (SLE6). Each dot indicates positive CosMx signal color coded by gene. **(B)** Heatmap of neighborhood-level ISG module activity indicated control and cLN tissue sections. White circles indicate location of glomeruli.

**Supplemental Figure 4:**
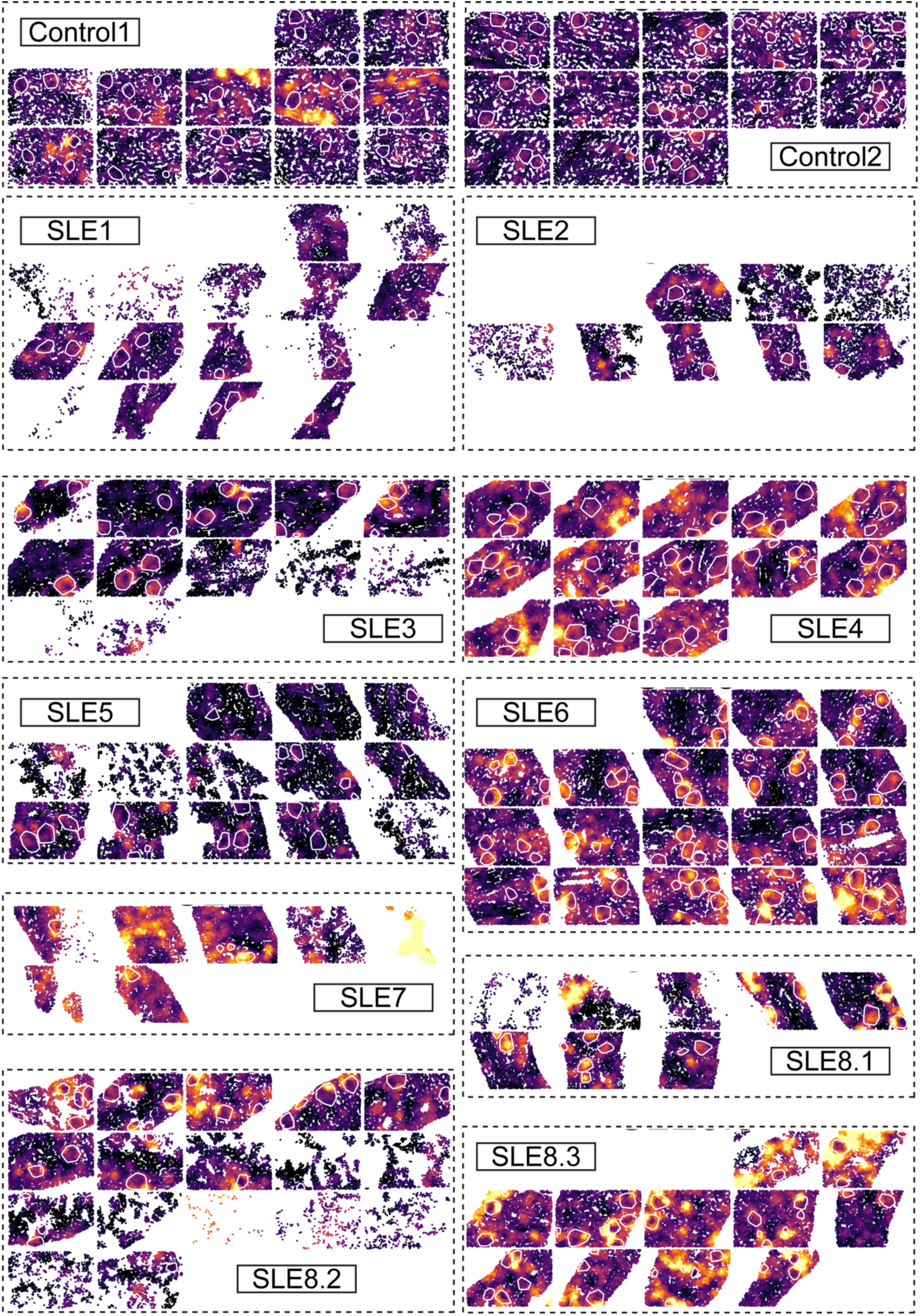
Macrophage “Class II HLA” spatial module in cLN. Heatmap of neighborhood-level “Class II HLA” module activity indicated control and cLN tissue sections. White circles indicate location of glomeruli.

**Supplemental Figure 5:**
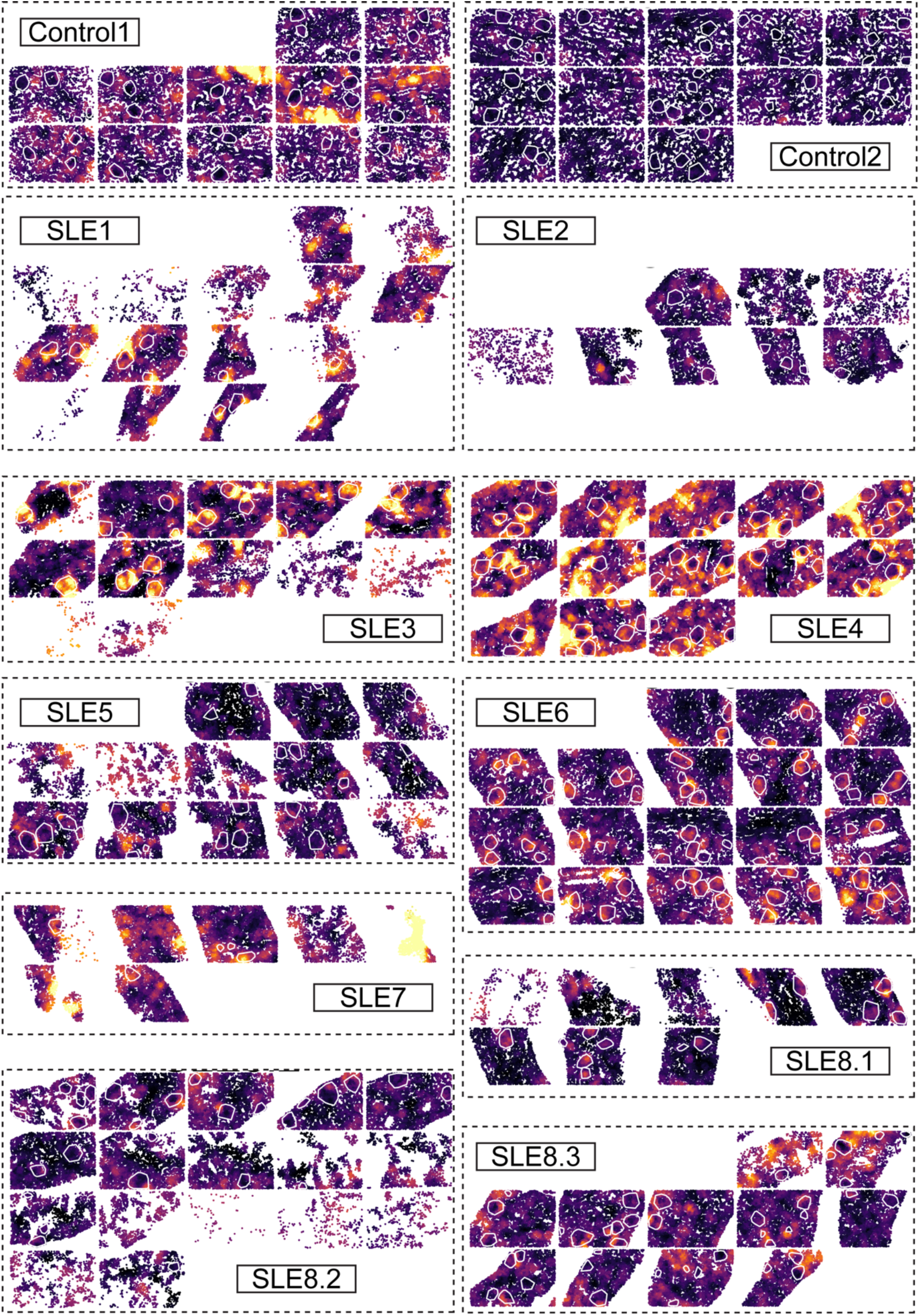
Macrophage “Complement” spatial module in cLN. Heatmap of neighborhood-level “Complement” module activity indicated control and cLN tissue sections. White circles indicate location of glomeruli.

**Supplemental Figure 6:**
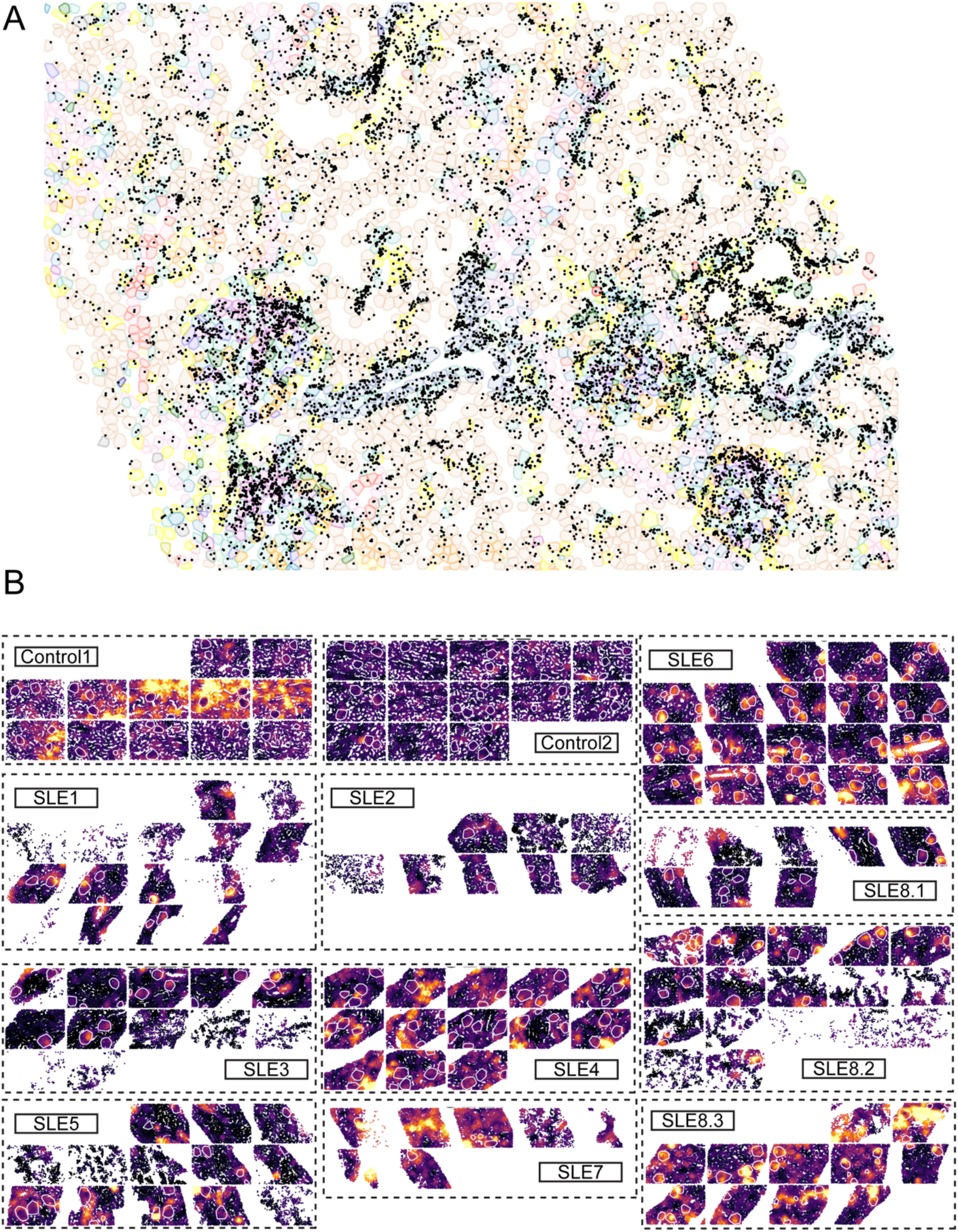
Fibrosis spatial module in cLN. **(A)** mRNA locations of genes within the “fibrosis” gene module in representative cLN kidney tissue (SLE6). Each black dot indicates expression of an individual module gene. **(B)** Heatmap of neighborhood-level “fibrosis” module activity indicated control and cLN tissue sections. White circles indicate location of glomeruli.

**Supplemental Table 1:** Patient demographics and clinical data in the cLN cohort.

| Patient | SLE1 | SLE2 | SLE3 | SLE4 | SLE5 | SLE6 | SLE7 | SLE8 |
| --- | --- | --- | --- | --- | --- | --- | --- | --- |
| Age (SLE diagnosis) | 12 years | 16 years | 11 years | 10 years | 15 years | 13 years | 13 years | 13 years |
| Age (kidney biopsy) | 12 years | 16 years | 11 years | 10 years | 15 years | 13 years | 13 years | 16, 17, 18 years |
| Sex | M | M | F | F | F | F | M | F |
| Race | Asian | Asian | White | Black / White | White | Other | White | White |
| Ethnicity (Hispanic) | No | No | Yes | No | No | Yes | No | Yes |
| Lupus Nephritis ISN/RPS Class | 3 | 3 | 4 | 3 | 3 | 4 | 4 | 4, 4, 3 |
| Extra-renal manifestations | Myositis, arthritis, malar rash, diarrhea, nasal ulceration, headaches, constitutional (fatigue, weight loss, fever) | Pleuritis, syncope, constitutional (weight loss, fatigue, fevers) | Arthritis, rash, fever, hepatitis | Alopecia, malar rash, palatal ulcer, arthritis | Necrotic abdominal lymph nodes, pancreatitis, rash, tachycardia, hypotension, hepatitis, Macrophage Activation Syndrome (MAS), constitutional (fever, weight loss) | Malar rash, rash on lower extremity | Arthritis, malar rash, hemolytic anemia, abdominal pain (vasculitis vs mesenteric adenitis) | Arthritis, rash |
| ANA (indirect immunofluorescence) | >1:640 | 1:160 | >1:5120 | >1:1280 | >1:640 | >1:320 | >1:320 | ANA positive (unknown titer) |
| Anti-dsDNA | Positive | Positive | Positive | Positive | Positive | Positive | Positive | Positive |
| Other serologies | RNP, Sm, Sm/RNP | Lupus inhibitor | Direct antiglobulin (Coombs) test (DAT) | RNP, Sm, Sm/RNP, MPO | SSA, SSB | | Anti- β2 glycoprotein IgG and IgM, Anti-cardiolipin IgG and IgM | Unknown |

**Supplemental Table 2:** Control subjects patient demographics and clinical data.

| **Patient** | **Control1** | **Control2** |
| --- | --- | --- |
| **Age (at nephrectomy)** | 17 years | 15 years |
| **Sex** | F | F |
| **Race** | White | White |
| **Ethnicity (Hispanic)** | Yes | No |
| **Diagnosis** | Nutcracker syndrome s/p left renal vein transposition. Subsequently, developed external left renal vein occlusion managed with failed renal autotransplant, and ultimately left nephrectomy. | Papillary Renal Cell Carcinoma, Type 1 |
| **Histopathology** | Essentially normal kidney parenchyma, except for reactive changes associated with renal vessels and renal serosa. | Kidney parenchyma distant from the tumor: glomeruli and tubules are viable, without sclerosis, inflammation, or interstitial fibrosis. |
| **Creatinine (at nephrectomy)** | 0.5 | 0.6 |
| **Estimated GFR (at nephrectomy)** | 135 | 114 |
| **Urinalysis (at nephrectomy)** | Negative for blood and protein | Negative for blood and protein |
| **Urine PCR (at nephrectomy)** | Not obtained | <0.2 |

**Supplemental Table 3:** Lupus nephritis immunosuppressive treatments (SLE1-SLE7)

| Immunosuppression prior to kidney biopsy |  |  |  | cLN treatment |  |  |  |  |  |  |
| --- | --- | --- | --- | --- | --- | --- | --- | --- | --- | --- |
| Patient | HCQ | Steroids | CYC |  | HCQ | Steroids | CYC | MMF | Belimumab | Tacrolimus |
| SLE1 | - | MP (1g x 3) | - |  | 300mg | Pred (30mg BID) | - | 2000 mg/day | - | - |
| SLE2 | 300mg | MP (1g x 3) | - |  | 300mg | MP (1g x 1), Pred (60mg qday) | - | 2000 mg/day | - | - |
| SLE3 | 200mg | MP (1g x 3), Pred (30mg BID) | - |  | 200mg | Pred (30mg BID) | 980mg x 1 | 2000 mg/day | - | - |
| SLE4 | - | MP (1g x 3), Pred (30mg BID) | - |  | 200mg | MP (1g x 1), Pred (30mg BID) | - | 1500 mg/day | + (for active arthritis / rash) | - |
| SLE5 | 400mg | MP (1g x 3), Pred (30mg BID) | 1 x dose (10 days prior to Bx) |  | 400mg | Pred (30mg BID) | 500mg x 1 | 2000 mg/day | - | - |
| SLE6 | - | Pred (20-80mg/day) | - |  | 200mg | Pred (80mg qday) | - | 2000 mg/day | - | - |
| SLE7 | - | - | - |  | 300mg | MP (1g x 1), Pred (60mg qday) | - | 1500 mg/day | - | + |

**Abbreviations:** HCQ, hydroxychloroquine; MP, methylprednisolone (IV); Pred, prednisone (oral); CYC, cyclophosphamide (IV); MMF, Mycophenolate Mofetil (oral)

**Supplemental Table 4:** Trend in laboratory values for 12 months after cLN diagnosis (SLE1-SLE7)

|  | Time of biopsy |  |  |  |  |  |  |
| --- | --- | --- | --- | --- | --- | --- | --- |
|  | Creatinine<br>(mg/dL) | eGFR<br>(ml/min/1.73m<sup>2</sup>) | Urine PCR<br>(mg/mg) | Albumin<br>(mg/dL) | Anti-dsDNA<br>(IU/mL; n<10) | Complement<br>C3 (mg/dL) | Complement C4<br>(mg/dL) |
| SLE1 | 0.8 | 90 | 1.0 | 3.5 | >300 | 34 | <7 |
| SLE2 | 0.9 | 81 | 1.1 | 3.4 | 55 | 107 | 44 |
| SLE3 | 0.7 | 90 | 4.3 | 2.5 | >300 | <22 | <7 |
| SLE4 | 0.5 | 118 | 5.8 | 2.3 | >300 | 32 | 7 |
| SLE5 | 0.4 | 174 | 2.2 | 2.2 | 306 | 32 | <7 |
| SLE6 | 0.8 | 80 | 5.3 | 1.8 | 70 | 42 | <8 |
| SLE7 | 0.9 | 83 | 16.7 | 1.6 | 175 | 40 | <7 |

|  | 6 Months |  |  |  |  |  |  |
| --- | --- | --- | --- | --- | --- | --- | --- |
|  | Creatinine<br>(mg/dL) |  | Urine PCR<br>(mg/mg) | Albumin<br>(mg/dL) | Anti-dsDNA<br>(IU/mL; n<10) | Complement<br>C3 (mg/dL) | Complement<br>C4 (mg/dL) |
| SLE1 | 0.7 | 104 | <0.2 | 4.6 | 7 | 108 | 17 |
| SLE2 | 0.8 | 91 | <0.2 | 4 | 6 | 106 | 30 |
| SLE3 | 0.5 | 126 | <0.2 | 5.2 | 16 | 117 | 19 |
| SLE4 | 0.3 | 197 | <0.2 | 4.6 | 52 | 119 | 21 |
| SLE5 | 0.7 | 100 | <0.2 | Unavailable | 6 (normal <4) | 160 | 22 |
| SLE6 | 0.5 | 128 | <0.2 | 3.6 | 18 | 112 | 18 |
| SLE7 | 0.6 | 126 | 0.52 | 3.5 | 5 | 96 | 14 |

|  | 12 months |  |  |  |  |  |  |
| --- | --- | --- | --- | --- | --- | --- | --- |
|  | Creatinine<br>(mg/dL) |  | Urine PCR<br>(mg/mg) | Albumin<br>(mg/dL) | Anti-dsDNA<br>(IU/mL; n<10) | Complement<br>C3 (mg/dL) | Complement C4<br>(mg/dL) |
| SLE1 | 0.7 | 105 | <0.2 | 4.5 | 3 | 107 | 18 |
| SLE2 | 0.9 | 81 | 0.2 | 4.7 | 4 | 117 | 28 |
| SLE3 | 0.3 | 213 | <0.2 | 5 | 9 | 110 | 17 |
| SLE4 | 0.3 | 200 | <0.2 | 4.6 | 29 | 141 | 30 |
| SLE5 | 0.7 | 100 | <0.2 | Unavailable | 4 | 135 | 21 |
| SLE6 | Unavailable | Unavailable | Unavailable | Unavailable | Unavailable | Unavailable | Unavailable |
| SLE7 | 0.8 | 96 | <0.2 | 4.1 | 4 | 102 | 14 |

**Supplemental Table 5:** B cell subcluster marker genes Genes were identified using the Seurat package and p-values adjusted using the Benjamini-Hochberg procedure.

| Gene | Subcluster | P value | avg_log2FC | P value adj |
| --- | --- | --- | --- | --- |
| CD37 | B.3 | 2.61E-20 | 0.923331 | 1.30E-18 |
| CD38 | B.4 | 9.54E-16 | 1.721163 | 4.77E-14 |
| CD52 | B.3 | 2.15E-31 | 4.48564 | 1.07E-29 |
| CD53 | B.3 | 5.87E-36 | 23.20923 | 2.93E-34 |
| CD79A | B.3 | 0.0091 | 0.386168 | 0.455007 |
| CIITA | B.3 | 7.62E-18 | 3.444508 | 3.81E-16 |
| CSK | B.3 | 0.004224 | 1.222816 | 0.211183 |
| CXCR4 | B.1 | 4.18E-08 | 0.41476 | 2.09E-06 |
| CXCR4 | B.3 | 0.000414 | 1.236 | 0.020721 |
| FKBP11 | B.4 | 4.75E-06 | 1.401699 | 0.000237 |
| HLA-DPA1 | B.3 | 5.25E-39 | 13.91653 | 2.62E-37 |
| HLA-DPB1 | B.3 | 5.75E-45 | 4.215574 | 2.88E-43 |
| HLA-DQA1 | B.3 | 6.71E-46 | 13.74758 | 3.35E-44 |
| HLA-DQB1 | B.2 | 0.001989 | 0.777247 | 0.099455 |
| HLA-DQB1 | B.3 | 6.22E-27 | 1.964848 | 3.11E-25 |
| HLA-DRA | B.3 | 4.03E-58 | 16.25719 | 2.01E-56 |
| HSP90B1 | B.1 | 1.33E-06 | 28.77536 | 6.63E-05 |
| IGHA1 | B.2 | 7.42E-107 | 52.15276 | 3.71E-105 |
| IGHD | B.1 | 4.85E-13 | 88.64234 | 2.42E-11 |
| IGHG1 | B.4 | 1.46E-135 | 54.78507 | 7.29E-134 |
| IGHG2 | B.4 | 6.19E-141 | 29.97054 | 3.10E-139 |
| IGHM | B.1 | 1.16E-105 | 88.24254 | 5.79E-104 |
| IGKC | B.3 | 2.68E-28 | 54.50807 | 1.34E-26 |
| IL16 | B.3 | 4.32E-21 | 1.588218 | 2.16E-19 |
| JCHAIN | B.1 | 1.45E-20 | 4.405219 | 7.27E-19 |
| JCHAIN | B.2 | 1.50E-40 | 1.915866 | 7.50E-39 |
| LAIR1 | B.3 | 0.001989 | 17.73385 | 0.099474 |
| LPAR5 | B.4 | 1.81E-29 | 2.246763 | 9.03E-28 |
| LYZ | B.3 | 1.01E-31 | 8.25153 | 5.03E-30 |
| MPO | B.3 | 0.000431 | 6.762452 | 0.021552 |
| MZB1 | B.4 | 1.62E-27 | 12.3535 | 8.09E-26 |
| PTPRCAP | B.3 | 1.08E-18 | 0.856019 | 5.39E-17 |
| ST6GAL1 | B.4 | 0.003854 | 2.30821 | 0.192712 |
| TNFRSF17 | B.3 | 1.16E-12 | 22.09005 | 5.80E-11 |
| XBP1 | B.4 | 2.66E-44 | 9.568072 | 1.33E-42 |

